# Comparison of expression profiling through microarray and RNA-seq analysis for Nipah virus

**DOI:** 10.1101/536284

**Authors:** Akanksha Rajput, Manoj Kumar

## Abstract

The Nipah virus is responsible various outbreaks among countries of south east Asia, most recent is in Kerala, India. It is considered to be highly contagious and having a range of vectors for transmission. The condition worsens due to the lack of effective inhibitors. This study is first study, which focused to detect the differentially expressed genes among two different NiV studies from 2012 and 2017. The transcriptomic profiling data were retrieved from the sequence archives. The multivariate gene enrichment analyses were performed on the log transformed data from them using pathway, gene ontology, disease, reactome, etc. The comparison study suggests that the down regulated differentially expressed genes are common among them as compared to up regulated ones with statistical significance. However, among the diseased category the upregulated genes are mostly from metabolic pathways and diseased category like metabolic pathways, heart failure, cholesterol metabolism while the downregulated genes linked to various cancers, and viral diseases like hepatitis, dengue, influenza, etc. We found various small molecules mapped in the pathways which are differentially expressed among the studies, which could be targeted so as to control the Nipah infection. In order to design the inhibitors, our study would be useful to extract the effective and broad-spectrum drug targets.

## Introduction

Nipah Virus (NiV) is a zoonotic virus from *paramyxoviridae* family and causes outbreaks in various countries of south-east Asia like Malaysia (1998-1999), Bangladesh (2001, 2003, 2004, 2005, 2007, 2008, 2009, 2010, 2011, 2012), Singapore (1999), India (2001, 2007, 2018) (http://www.searo.who.int/entity/emerging_diseases/links/nipah_virus_outbreaks_sear/en/). Its name “Nipah” comes from the Malaysian village name “Sungai Nipah” that causes encephalitis in pig farmers. The morbidity and mortality ranges from 0-100% among various outbreaks. Interestingly, most of the outbreaks are limited by seasons and geographical range e.g. between winter and spring and during breeding season of bats and date palm sap harvesting season. It is transmitted to humans from infected bats, pigs, and humans (https://www.cdc.gov/vhf/nipah/). The *Paramyxoviridae* family viruses Nipah and Hendra virus possess similar genome organization (Harcourt et al., 2000).

NiV is negative single stranded RNA (ssRNA) virus, with ∼18kb genome size with six transcriptional units encoding nine proteins. The nine proteins are nucleocapsid (N), phosphoprotein (P), fusion protein (F), glycoprotein (G), polymerase (L), matrix protein (M), and interferon antagonists W, V and C (Martinez-Gil et al., 2017). The unique characteristics of the members of the genus *Henipavirus* from other paramyxovirus is the presence of long untranslated regions (UTR) at 3’ end of viral mRNA transcripts excluding L gene (Harcourt et al., 2000). Among the important proteins the phosphoprotein (P) is a polymerase cofactor and enhance the polymerase activity as well as allows the encapsulation of newly synthesized viral genomes and antigenomes (Yoneda et al., 2010). However, the glycoprotein G helps the virus to attach with the host cell through ephrin B2 and B3 (Negrete et al., 2005). Thus, every gene has a vital role to play in NiV life cycle (Jensen et al., 2018). During the infection of NiV in the host, it modifies (up/down) the expression of various genes within the host. The change of dynamics of genes in host cell is an important tool to explore viral biology including its pathogenesis.

The gene expression-based profiling is an important and highly sensitive tool to measure the activity of genes at a time. Various transcriptomic technologies like microarrays, RNAseq, DNAseq, CHIPseq, etc have been used to generate and analyze the data (Bucca et al., 2004). Microarray is beneficial technology to differentiate the gene expression between two mRNA samples (Govindarajan et al., 2012). In general, it is used to identify the set of genes over- or under-expressed between diseased and normal cell. While the RNAseq based expression profile is more advantageous to detect the up or down regulated genes from known as well as new transcripts with more sensitivity (Wang et al., 2009). Hence, the gene expression profiling would be highly advantageous to investigate the NiV pathogenesis.

In the present study we are utilizing the gene expression-based profiling to extract the differentially expressed genes (DEGs) from NiV two studies from year 2012 and 2017 (Mathieu et al., 2012; Martinez-Gil et al., 2017). As NiV is a BSL4 pathogen, therefore working on live virus is very limited. In literature the limited amount of computational analyses (intra-study) was performed to investigate the modified genes during infection (Mathieu et al., 2012; Martinez-Gil et al., 2017). However, the comprehensive cross comparison study between the two studies has not been done in the literature, so as to detect the common DEGs between the NiV infection. Thus, in current study we explored both intra and inter studies DEGs to investigate the pathogenesis of NiV. We performed extensive computational analysis for the DEGs involved in pathways; disease condition; Gene ontology (GO) based on molecular function, biological process, and cellular component; reactomes; and many more.

## Material and Methods

### Data collection

For analyzing the gene expression profiling of the NiV, the data were extracted from NCBI Gene Expression Omnibus (GEO) and Sequence Read Archive (SRA) database. In 2012 Mathieu *et al* performed microarray experiment of NiV isolate UMMC1 infection (Malaysian strain) of primary endothelial cells i.e. HUVEC between infected and unfected cells (Mathieu et al., 2012). In 2017 Martinez-Gil *et al* performed RNAseq experiment on NiV infected and uninfected HEK293T cell line (Martinez-Gil et al., 2017). The sample id of mock and experiment of the 2012 were GSM813064, GSM813066 and GSM813065, GSM813067 while that of 2017 were SRS2461913, SRS2461912, SRS2461909 and SRS2461911, SRS2461910, SRS2461908 respectively. Further, the DEG were extracted from the studies and subjected to various downstream analysis on both intra and inter studies.

### Data preprocessing

Microarray and RNAseq data were preprocessed for the downstream processes. The differentially expressed genes were extracted from the samples through alignment, assembly and normalization of the data. The reads of microarray data were processed using limma package of Bioconductor (Ritchie et al., 2015). However, the RNA-seq data was refined using tophat2/cufflink/cuffdiff package (Trapnell et al., 2012). For the DEGs log fold change value of ±1.5 was used for further analysis. For microarray 298 up and 341 down regulated genes, while for the RNAseq 259 up and 1664 down regulated were extracted using above mentioned strategies.

### Pathway analysis

The pathway analysis of the up and down regulated genes was accomplished to check the distribution of genes among important pathways of humans using Kyoto Encyclopedia of Genes and Genomes (KEGG)(Kanehisa et al., 2017). The KEGG module in DAVID software extracts the molecular interactions, relations and reactions among the genes and resulted in the significant pathways in which they are involved.

### Gene ontology analysis

Gene ontology enrichment analysis helps in the identification of group of genes, which possess common molecular functions, biological processes, and cellular component. We performed GO based analysis through Gene Ontology Consortium (Ashburner et al., 2000). The up and down regulated genes from both studies were evaluated for all the three domains of GO.

### Disease category

The DEGs were explored for the diseased category using Generalized anxiety disorder (GAD) for checking the anxiety disorder. To check the disorders, we used DAVID software for the gene enrichment analysis (Huang da et al., 2009). The significantly up and down regulated from both the studies were used for intra and inter studies analysis.

### Reactome

We used reactome pathway software to map the pathways, macromolecular complexes, reactions (Croft et al., 2011). It helps to explore all the gene enrichment analysis for up and downregulated genes among both the studies.

### RRHO

We have used Rank-Rank Hypergeometric Overlap (RRHO) for comparing the genes between two studies. The RRHO is used to rank the genes according to their effect size and *p*-values according to the degree of differential expression (Plaisier et al., 2010). We constructed Rank scatter plot and RRHO map to compare the 2012 and 2017 NiV studies. The R package for RRHO was used for the same available in Bioconductor.

## Result

### Pathway analysis

The upregulated genes of 2012 Malaysian strain showed to be existed maximally in Metabolic pathways with 12 genes followed by Neuroactive ligand-receptor interaction, Cytokine-cytokine receptor interaction, Calcium signaling pathway, cGMP-PKG signaling pathway, Purine metabolism, Chemokine signaling pathway having 10, 7, 4, 4, 3 and 3 genes respectively (**Figure 1A**). The downregulated genes reported to be existed in Influenza A, Hepatitis C, Measles, Herpes simplex infection, Cytokine-cytokine receptor interaction, Metabolic pathways, Cytokine-cytokine receptor interaction with 15, 12, 12, 12, 11, 10, 10 correspondingly (**Supplementary Figure 1A**).

**Figure 1.**
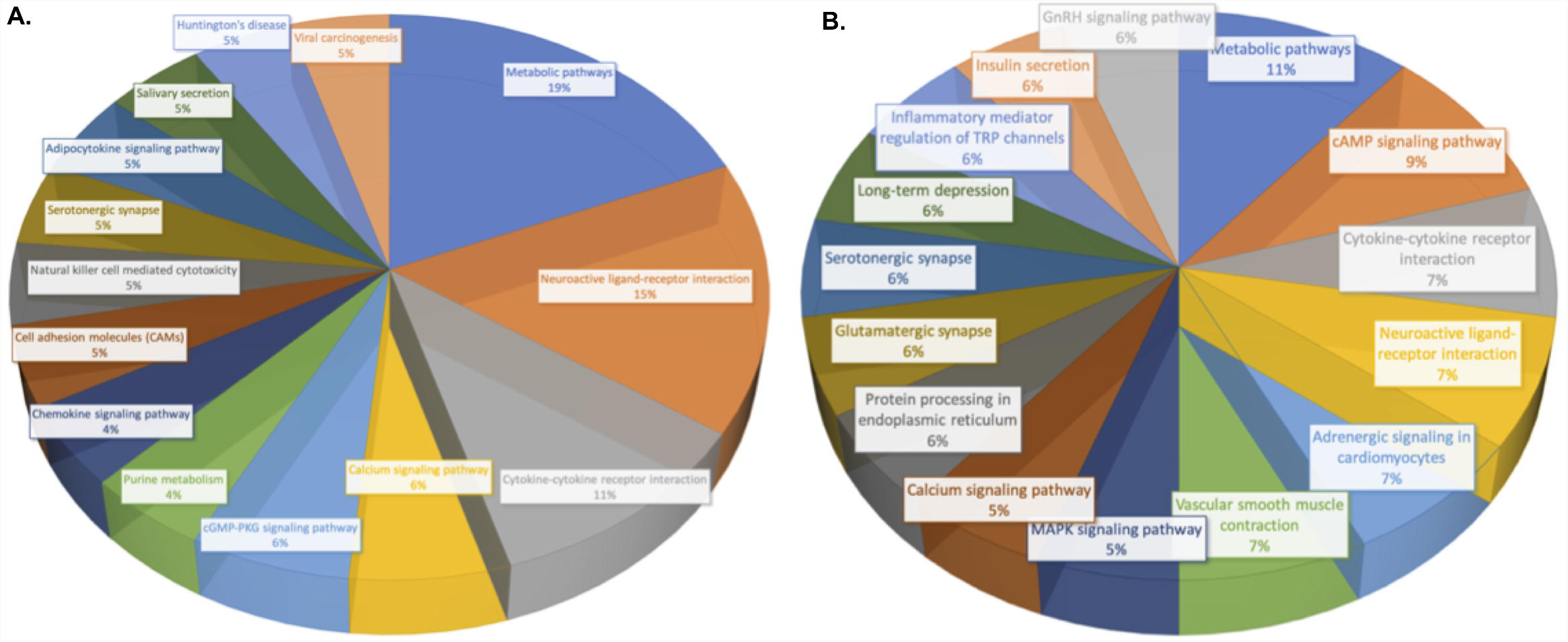
Pie chart depicting differentially expressed upregulated genes having pathway mapped **A.** 2012 study **B.** 2017 study

The 2017 strain of NiV showed that maximum upregulated genes lies in pathway of Metabolic pathways, cAMP signaling pathway, Cytokine-cytokine receptor interaction, Neuroactive ligand-receptor interaction, Adrenergic signaling in cardiomyocytes, Vascular smooth muscle contraction with 6, 5, 4, 4, 4, 4 respectively (**Figure 1B**). The downregulated genes of 2017 strain seems to be mapped in Alcoholism, Systemetic lupus erythematosus, Metabolic pathways, Cytokine-cytokine receptor interaction, Neuroactive ligand-receptor interaction, Viral carcinogenesis, PI3K-Akt signaling pathways with 46, 46, 40, 37, 27, 24, 20 genes correspondingly (**Supplementary Figure 1B**).

Interestingly, both the studies of NiV showed that upregulated genes have common pathways in Cytokine-cytokine receptor interaction, Neuroactive ligand-receptor interaction, Metabolic pathways, and Calcium signaling pathways. While downregulated genes mapped in PI3K-Akt signaling pathway, Influenza A, Measles, Tuberculosis, Jak-STAT signaling pathway, etc (**Supplementary Figure S2**).

### Gene ontology analysis

The 2012 study shows that upregulated genes have the molecular function protein binding; metal ion binding; DNA binding; ATP binding; Calcium ion binding; Zinc ion binding; Identical protein binding with 83; 18; 15; 13; 10; 10; 10 (**Supplementary Figure S3A**). While the downregulated genes lie in the topmost molecular function protein binding; metal ion binding; DNA binding; ATP binding; zinc ion binding; transcription factor activity; G-protein coupled receptor activity with 93; 23; 20; 20; 19; 14; 14 respectively (**Supplementary Figure S4A**). The biological process with upregulated genes is transcription, DNA-templated; signal transduction; positive regulation of transcription from RNA polymerase II; regulation of transcription, DNA-templated; spermatogenesis; Immune response; Cell adhesion with 16; 16; 12; 11; 11; 10; 10 (**Figure 2A**). The downregulated genes involved defense response to virus; signal transduction; transcription, DNA-templated; innate immune response; positive regulation of transcription from RNA polymerase II; type I interferon signaling pathway with 24; 23; 21; 19; 19 correspondingly (**Supplementary Figure S5A**). The cellular component of the upregulated genes followed in integral component of membrane; plasma membrane; nucleus; cytoplasm; integral component of plasma membrane; extracellular exosome; extracellular space with 64; 57; 46; 45; 33; 32; 30 genes (**Supplementary Figure S6A**) while downregulated genes lies in cellular component like cytoplasm; integral component of membrane; plasma membrane; nucleus; cytosol; extracellular region with 68; 68; 59; 58; 42 and 32 genes (**Supplementary Figure S7B**).

**Figure 2.**
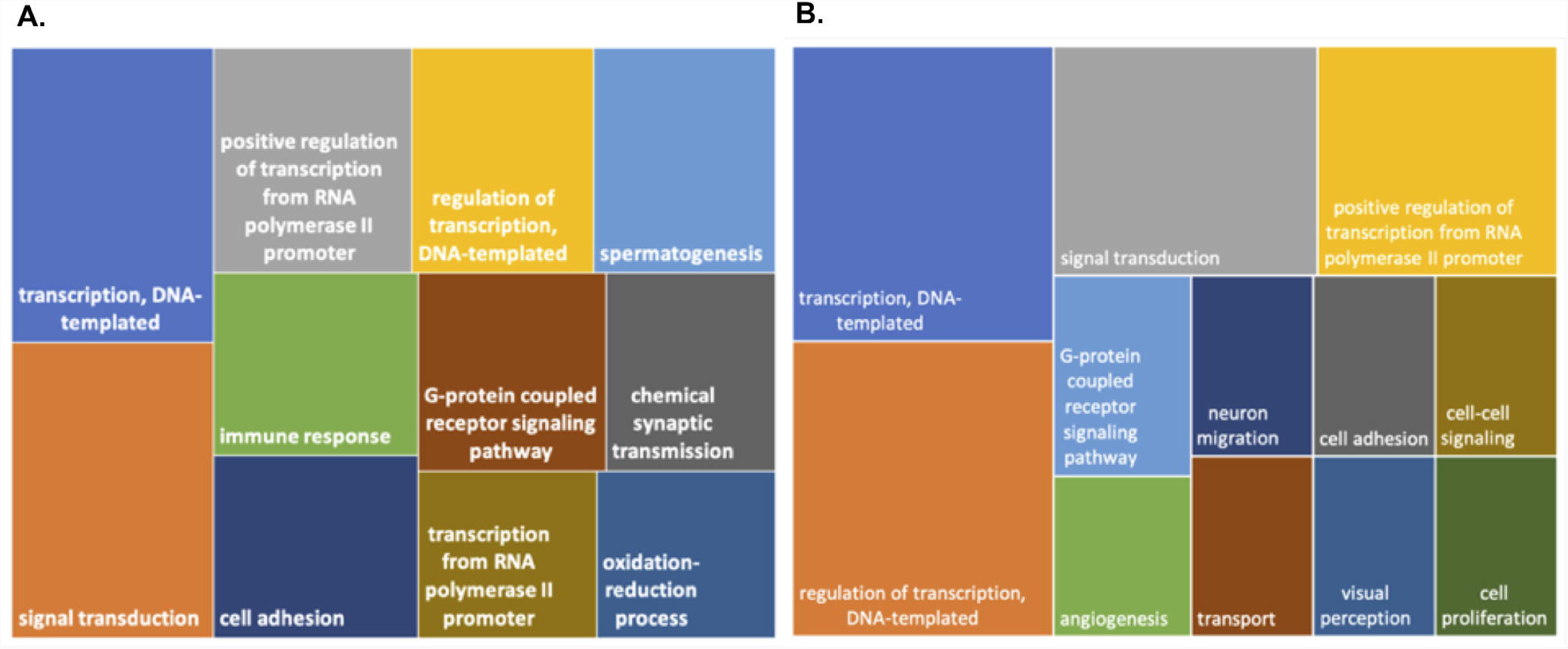
Tree Map depicting differentially expressed upregulated genes having gene ontology based biological process from **A.** 2012 study **B.** 2017 study

The 2017 study displays that upregulated genes possess the molecular functions like protein binding, metal ion binding, nucleic acid binding, zinc ion binding, DNA binding, ATP binding, receptor binding with 45, 13, 9, 9, 8, 8, 7 (**Supplementary Figure S3B**). Whereas, the dowregulated genes showed the molecular functions like protein binding, DNA binding, protein heterodimeriation activity, metal ion binding, G-protein coupled receptor activity, ATP binding with 269, 60, 56, 49, 46, 41, 29 correspondingly (**Supplementary Figure S4B**). The upregulated genes with transcription, DNA-templated; regulation of transcription, DNA-templated; signal transduction; positive regulation of transcription from RNA polymerase II; G-protein coupled receptor signaling pathway; angiogenesis with 14; 14; 11; 10; 5; 4 correspondingly (**Figure 2A**). The downregulated genes mapped in pathway signal transduction; G-protein coupled receptor signaling pathway; transcription, DNA-templated; immune response; inflammatory response; nucleosome assembly with 65; 56; 40; 40; 39; 36 respectively (**Supplementary Figure S5B**). The upregulated genes having cellular component having nucleus; cytoplasm; integral component of membrane; plasma membrane; extracellular exosome; extracellular region; cytosol with 30; 28; 28; 26; 20; 18; 17 respectively (**Supplementary Figure S6B**). Whereas the downregulated gene having integral component of membrane; plasma membrane; nucleus; extracellular exosome; cytoplasm; extracellular region; extracellular space; integral component of plasma membrane having 261; 191; 172; 155; 143; 142; 123 and 90 genes correspondingly (**Supplementary Figure S7B**).

Among 2012 and 2017 NiV studies the common upregulated and downregulated genes with GO based molecular functions are 13 and 63 respectively and the details of all the intersecting genes are provided in **Supplementary Figure S8**. However, the common DEGs among GO based biological process are 14 and 253 correspondingly as shown in **Supplementary Figure S9**. While the GO based cellular component displayed the 2 and 27 up and downregulated genes as depicted in **Supplementary Figure S10**.

### Disease category

The 2012 study displayed that upregulated genes responsible to cause Myocardial infraction, Alcoholism, Psychiatric disorders, Obesity, Heart failure, Atherosclerosis with count of 12, 11, 10, 10, 10, 9 and *p*-value of 0.05, 0.07, 0.008, 0.03, 0.03, 0.05 (**Figure 3A**) while the downregulated genes falls in the category of Type 2 diabetes, Schizophrenia, Dengue Hemorrhagic fever, Multiple sclerosis, Hepatitis C, Diabetes Type 1 with 40, 17, 14, 14, 11, 10 gene count and *p*-value of 0.05, 0.07, 1.28E-09, 0.05, 3.11E-06, 0.0012 correspondingly (**Supplementary Figure S11A**).

**Figure 3.**
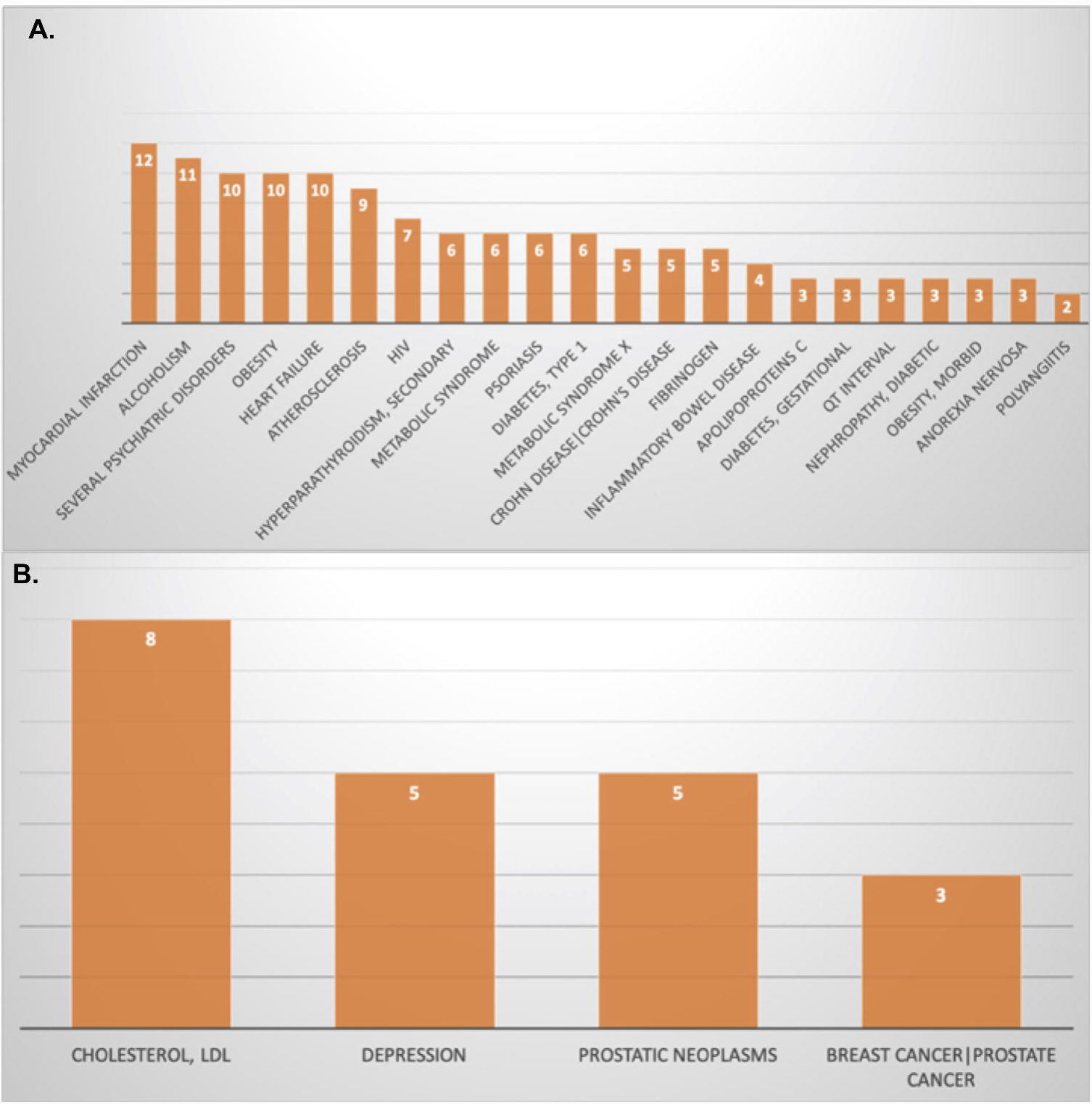
Bar graph showing the differentially expressed upregulated genes having GAD disease category from **A.** 2012 study **B.** 2017 study

The 2017 NiV study revealed that the upregulated genes mapped with the GAD disease like Cholestrol (LDL), Depression, Prostatic Neoplasms, Breast cancer/Prostate cancer with frequency of 8, 5, 5, 3 with *p*-value of 0.045, 0.02, 0.02, 0.06 (**Figure 3B**) the downregulated genes lies in the category of Type 2 diabetes, Chronic renal failure, Schizophrenia, Asthma, Hypertension, Lung Cancer with 120, 56, 42, 38, 37, 36 genes having p-value of 3.26E-04, 0.0014, 0.055, 5.07E-05, 0.09, 0.015 respectively (**Supplementary Figure S11B**).

### Reactome

The 2012 studies showed that upregulated genes were mapped in topmost pathways like signal transduction, metabolism, developmental biology with 29, 22 and 13 genes as shown in **Figure 4**. The signal transduction pathway effects the 08 gene involved in GPCR, secondary messengers, nuclear receptors, RHO GTPases as shown in **Supplementary Figure S12**. The metabolism pathway known to effect the genes involved in citric acid cycle, carbohydrate metabolism, porphyrin and lipid metabolism (**Supplementary Figure S13**). Moreover, the list of the molecules grouped the signal transduction and metabolism are listed in **Supplementary Tables S1 and S2** respectively. The components of the signal transduction pathway shown to map the chemicals like estrogen, estranol, retinyl, progesterone, etc., proteins like PDGFA, AGT, RHO, etc, whereas the metabolism pathway showed metal ions like Zn, Mg, amino acids like L-Glutamate, L-Proline, etc, proteins like RPL39L, ARF, etc. Downregulated gene displayed that among all the pathways as shown in **Supplementary Figure S14** most important pathways with maximum mapped genes are immune system, metabolism of protein, signal transduction with 76, 23 and 24 genes. The immune system pathways known to effect the components like cytokine, innate, and adaptive immune system as shown in **Supplementary Figure S15**. While the majority of the signal transduction pathway seems to be affected by the genes which are involved in signaling by nuclear receptors, WNT, Notch, MAPK, Tyrosine Kinase, secondary messengers, etc (**Supplementary Figure S16**). The small molecules like metals, heparan sulfate, L-Glutamate, ATP, etc., known to mapped in signal transduction pathways, whereas the trehalose, lipoteichoic acid, peptides, etc are for immune system as shown in **Supplementary Tables S3 and S4** respectively.

**Figure 4.**
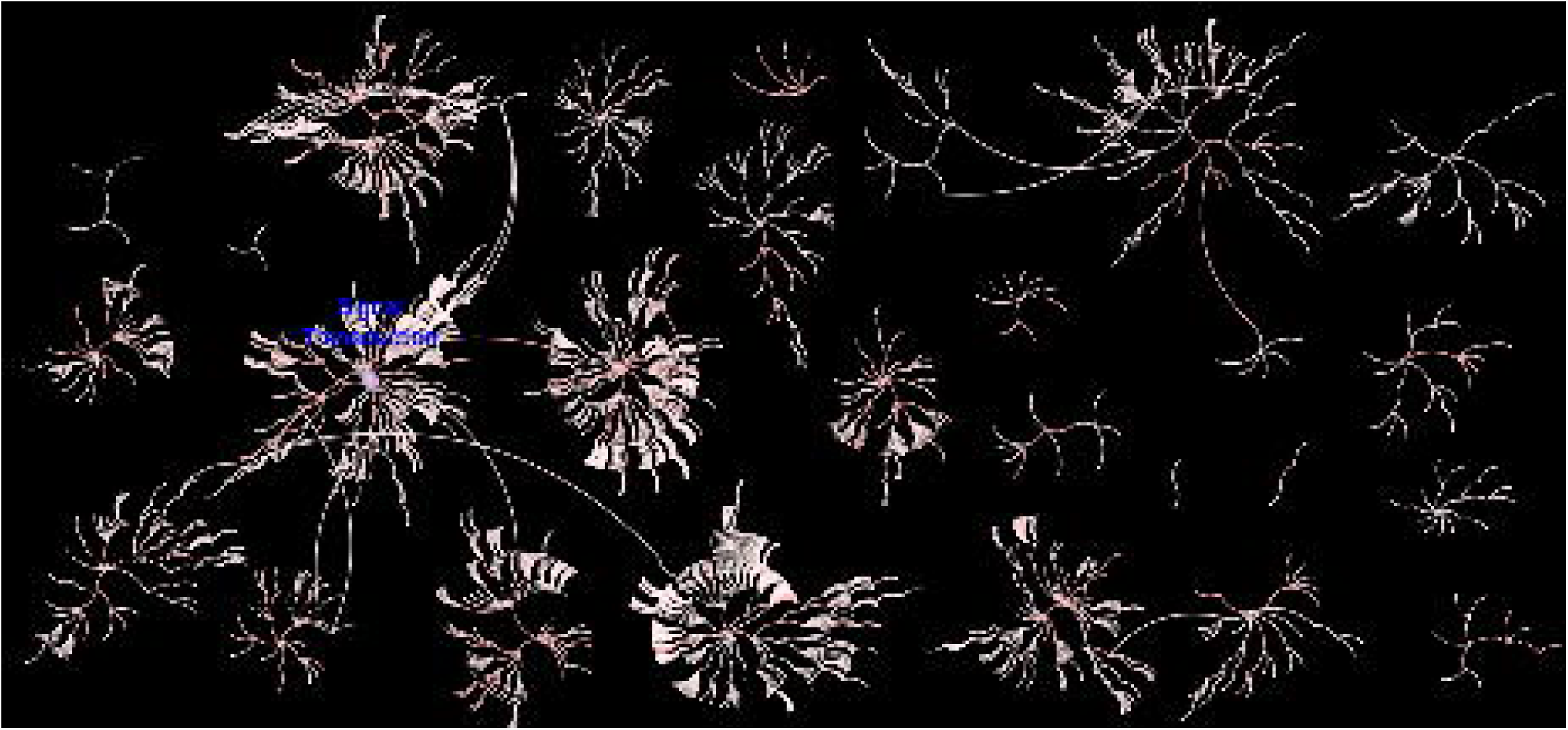
Pathway mapped of the upregulated genes from 2012 study using reactome software

In 2017 NiV study displayed that upregulated genes showed the topmost pathways like Metabolism, Signal transduction, Immune system, Transport of small molecules, with 18, 16, 8 and 8 genes as shown in **Figure 5**. The metabolism pathway components like amino acids metabolism, citric acid cycle, carbohydrate, nucleosides, etc., (**Supplementary Figure S17**) however, for the signal transduction pathway components effected mostly are RHO GTPases, secondary messengers, GPCR, WNT, and many more as shown in **Supplementary Figure S18**. However, the small molecules, proteins, DNA/RNA effecting both signal transduction and metabolism are enlisted in **Supplementary Tables S5 and S6** respectively. The downregulated genes mapped on the topmost pathways like signal transduction, Metabolism, gene expression, developmental biology with 140, 72, 67, 65 genes as shown in **Supplementary Figure S19**. The component of signal transduction pathway like TGF-Beta, Hedgehog, Notch, death receptor, WNT etc., (**Supplementary Figure S20**) while the metabolism pathways like Inositol phosphate, lipids, carbohydrates, nucleotides, are effected most (as shown in **Supplementary Figure S21**). However, the small chemicals, proteins, DNA/RNA mapped in the metabolism and signal transduction are enlisted in **Supplementary Tables S7 and S8** respectively.

**Figure 5.**
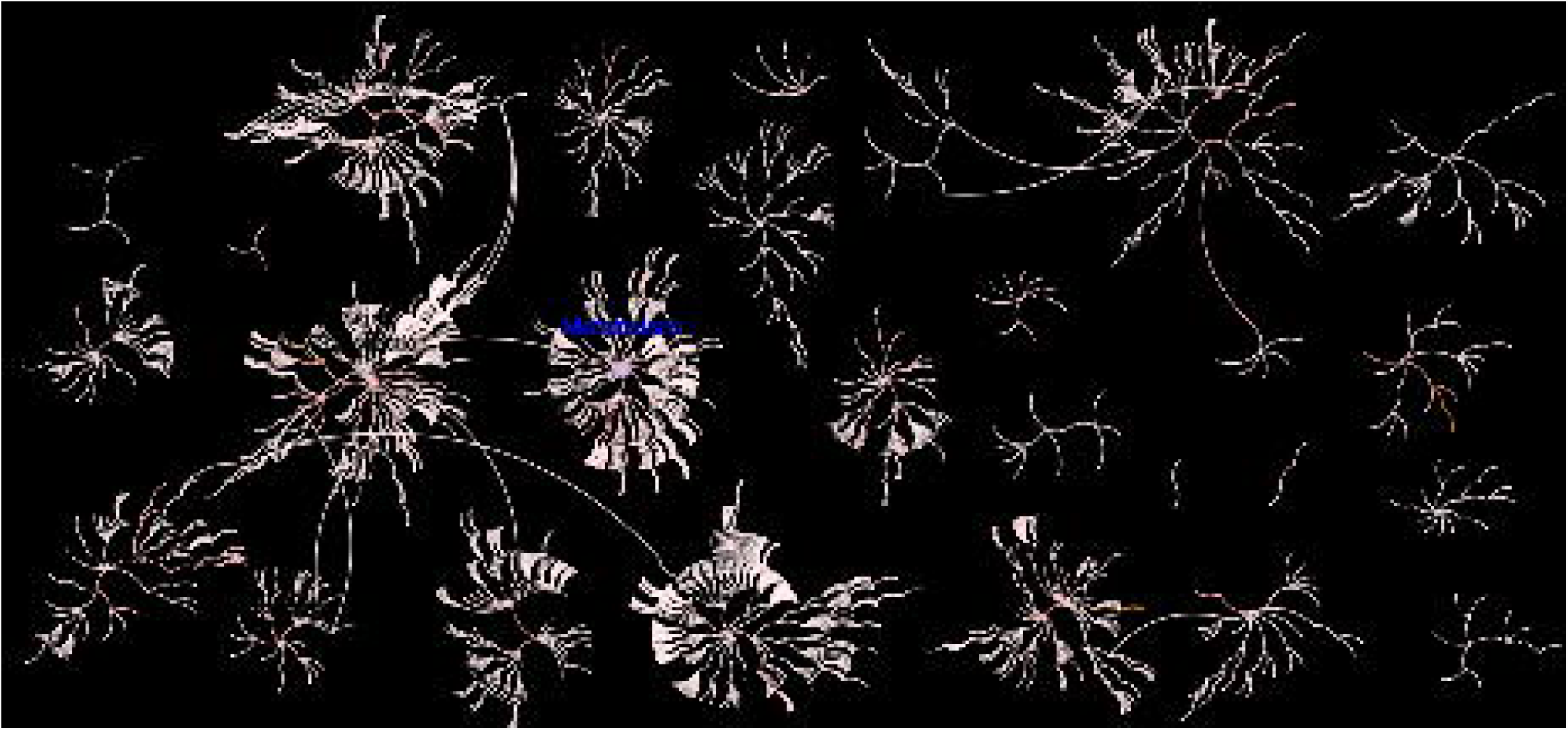
Pathway mapped of the upregulated genes from 2017 study using reactome software

### RRHO

The RRHO map depicts that the downregulated genes showed similarity between among each other followed by the upregulated genes with high statistical significance. However there is very less similarity is displayed between up and downregulated genes of both the studies and vice versa as shown in **Figure 6**. The gene ranking common in both the studies is explored by Rank-Rank scatter plot. The plot depicts that the gene expression profiles between both the studies were not so strongly correlated with spearman’s correlation coefficient of 0.0005 as shown in **Supplementary Figure S22**. The venn diagram of DEGs from both the studies are shown in **Supplementary Figure S23** with 221 upregulated and 824 upregulated genes.

**Figure 6.**
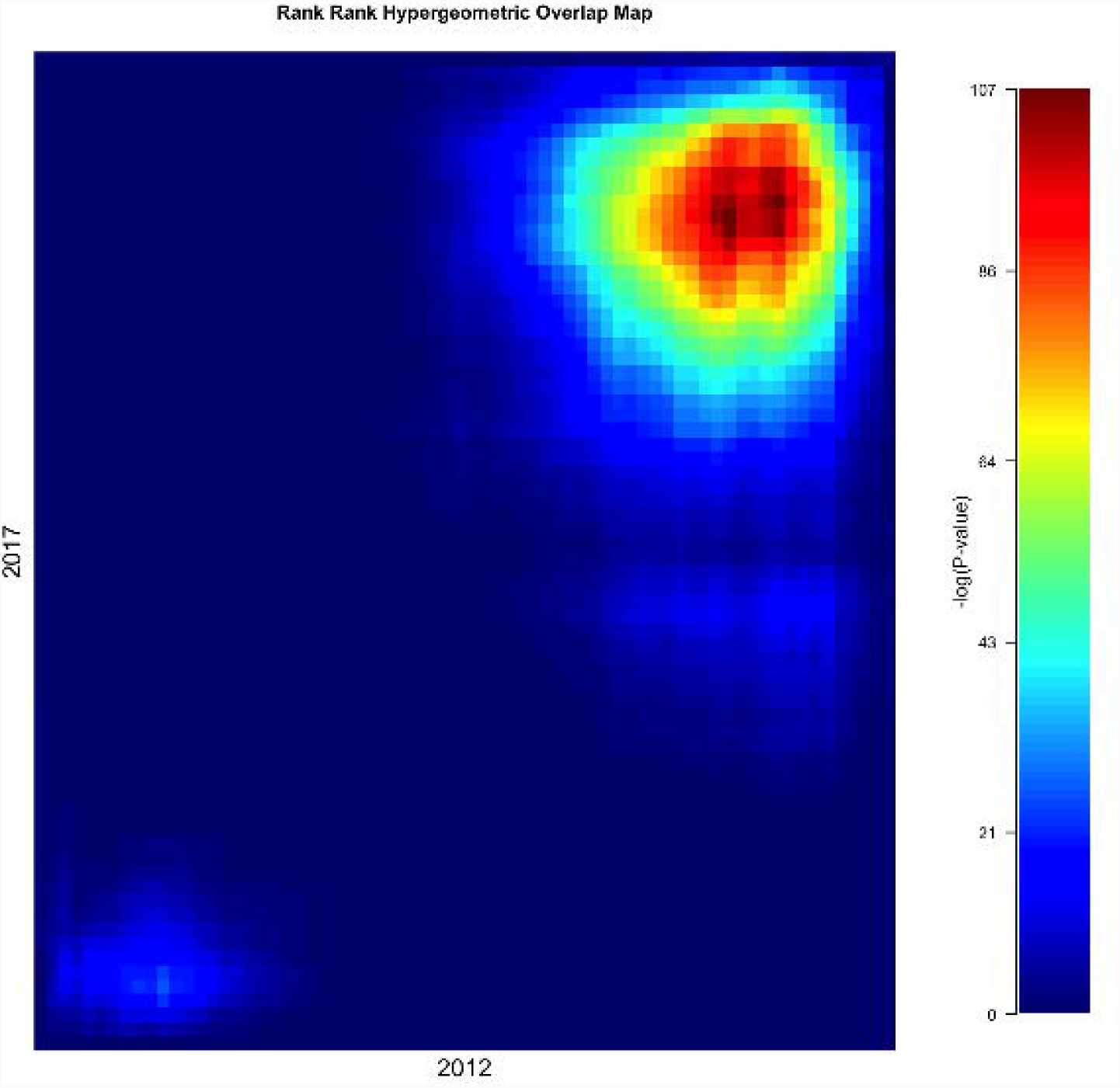
Rank-Rank Hypergeometric Overlap map showing the comparison between 2012 and 2017 studies

## Discussion

The NiV virus outbreaks is common in south eastern countries of Asia (Ang et al., 2018)(29643201). Its mortality varies as per the outbreak and till date there is no inhibitors of NiV due to the constraints of being BSL4 pathogen. In literature few studies have been done regarding the development of inhibitors (Dawes et al., 2018; Mathieu et al., 2018). In the current study, we identified the DEGs among two different studies and tried to find out the common and different genes, which are up and down regulated. Till date no study has been performed to analyze the DEGs against NiV. The multidimensional analyses would be helpful to find out the highly potent drug targets.

The pathway analysis showed that maximally modifies pathway during NiV infections are Metabolic pathways, Cytokine-cytokine receptor interaction, and Neuroactive ligand-receptor interaction. Previous studies showed that majority of the proteins effect chemokine signaling component with CXCL10 chemokine (Mathieu et al., 2012). However, our study showed that NiV infection alters various other important pathways in hosts that has not been reported till date.

GO based analysis showed that significant DEGs between two studies of NiV infection shows changes in specific type of pathways that are not extensively reported till date. The molecular functions like protein binding, metal ion binding (Zn, Ca), DNA binding, ATP binding. However, during the NiV infection the biological processes like transcription, signal transduction, G-protein coupled signaling pathway were altered significantly. The majorly changed cellular components are plasma membrane, cytoplasm and nucleus among significant DEGs. Thus, during the NiV infection the host pathways responsible for signaling and immune system effected commonly.

On scanning the GAD disease category for DEGs, we found that the upregulated genes for the metabolism, neurological, heart related showed more prominence, while the downregulated genes mostly fall in the category of other viral diseases, cancer and diabetes with good statistical significance. Thus, both the studies show intersection at some diseases.

The reactome analysis showed that in all the DEGs from both the studies maximally effect the critical pathways like metabolism, signal transduction, developmental biology, gene expression, etc. However, there are differences in the specific components among the common pathways that are changed during the intracellular and interstudies of Nipah virus. Therefore, we hypothesize that mode of action of NiV among both the studies are different.

While comparing the similarity between two studies we found that DEGs among both the studies were not so statistically similar among each other, while the downregulated genes were more similar as compared to upregulated genes. Therefore, both the NiV studies were not so similar in DEGs. Our study suggests that the most potent pathway to be targeted is signal transduction, specifically GPCR ligand binding of classA/1. However, most effective and specific targets common for both the studies are phospholipid phospahatase related 5 and adrenoceptor alpha 1D.

## Conclusion

Our study correlates the 2012 and 2017 NiV studies and examine the DEGs among them. This is the first study to compare microarray and RNAseq studies using log transformation of the fold change among them. We found that broad spectrum targets for the NiV are available, which can be used for developing effective anti-virals. For example, the pathways like metabolism, neurological disorder, cholesterol, heart disease would be targeted for the same because some genes among them seems to be upregulate during the NiV infection. Howeverm the current study also gives an insight for the drug repurposing against NiV. Therefore, we hope that this study would be helpful in extracting novel drug targets for NiV.

## Acknowledgements

We acknowledge the infrastructure support of Department of Biotechnology, Government of India (GAP0001).

## References

Ang, B.S.P., Lim, T.C.C., and Wang, L. (2018). Nipah Virus Infection. J Clin Microbiol 56(6). doi:10.1128/jcm.01875-17.

Ashburner, M., Ball, C.A., Blake, J.A., Botstein, D., Butler, H., Cherry, J.M., et al. (2000). Gene ontology: tool for the unification of biology. The Gene Ontology Consortium. Nat Genet 25(1), 25–29. doi:10.1038/75556.

Bucca, G., Carruba, G., Saetta, A., Muti, P., Castagnetta, L., and Smith, C.P. (2004). Gene expression profiling of human cancers. Ann N Y Acad Sci 1028, 28–37. doi:10.1196/annals.1322.003.

Croft, D., O’Kelly, G., Wu, G., Haw, R., Gillespie, M., Matthews, L., et al. (2011). Reactome: a database of reactions, pathways and biological processes. Nucleic Acids Res 39(Database issue), D691–697. doi:10.1093/nar/gkq1018.

Dawes, B.E., Kalveram, B., Ikegami, T., Juelich, T., Smith, J.K., Zhang, L., et al. (2018). Favipiravir (T-705) protects against Nipah virus infection in the hamster model. Sci Rep 8(1), 7604. doi:10.1038/s41598-018-25780-3.

Govindarajan, R., Duraiyan, J., Kaliyappan, K., and Palanisamy, M. (2012). Microarray and its applications. J Pharm Bioallied Sci 4(Suppl 2), S310–312. doi:10.4103/0975-7406.100283.

Harcourt, B.H., Tamin, A., Ksiazek, T.G., Rollin, P.E., Anderson, L.J., Bellini, W.J., et al. (2000). Molecular characterization of Nipah virus, a newly emergent paramyxovirus. Virology 271(2), 334–349. doi:10.1006/viro.2000.0340.

Huang da, W., Sherman, B.T., and Lempicki, R.A. (2009). Bioinformatics enrichment tools: paths toward the comprehensive functional analysis of large gene lists. Nucleic Acids Res 37(1), 1–13. doi:10.1093/nar/gkn923.

Jensen, K.S., Adams, R., Bennett, R.S., Bernbaum, J., Jahrling, P.B., and Holbrook, M.R. (2018). Development of a novel real-time polymerase chain reaction assay for the quantitative detection of Nipah virus replicative viral RNA. PLoS One 13(6), e0199534. doi:10.1371/journal.pone.0199534.

Kanehisa, M., Furumichi, M., Tanabe, M., Sato, Y., and Morishima, K. (2017). KEGG: new perspectives on genomes, pathways, diseases and drugs. Nucleic Acids Res 45(D1), D353– d361. doi:10.1093/nar/gkw1092.

Martinez-Gil, L., Vera-Velasco, N.M., and Mingarro, I. (2017). Exploring the Human-Nipah Virus Protein-Protein Interactome. J Virol 91(23). doi:10.1128/jvi.01461-17.

Mathieu, C., Guillaume, V., Sabine, A., Ong, K.C., Wong, K.T., Legras-Lachuer, C., et al. (2012). Lethal Nipah virus infection induces rapid overexpression of CXCL10. PLoS One 7(2), e32157. doi:10.1371/journal.pone.0032157.

Mathieu, C., Porotto, M., Figueira, T.N., Horvat, B., and Moscona, A. (2018). Fusion Inhibitory Lipopeptides Engineered for Prophylaxis of Nipah Virus in Primates. J Infect Dis 218(2), 218–227. doi:10.1093/infdis/jiy152.

Negrete, O.A., Levroney, E.L., Aguilar, H.C., Bertolotti-Ciarlet, A., Nazarian, R., Tajyar, S., et al. (2005). EphrinB2 is the entry receptor for Nipah virus, an emergent deadly paramyxovirus. Nature 436(7049), 401–405. doi:10.1038/nature03838.

Plaisier, S.B., Taschereau, R., Wong, J.A., and Graeber, T.G. (2010). Rank-rank hypergeometric overlap: identification of statistically significant overlap between gene-expression signatures. Nucleic Acids Res 38(17), e169. doi:10.1093/nar/gkq636.

Ritchie, M.E., Phipson, B., Wu, D., Hu, Y., Law, C.W., Shi, W., et al. (2015). limma powers differential expression analyses for RNA-sequencing and microarray studies. Nucleic Acids Res 43(7), e47. doi:10.1093/nar/gkv007.

Trapnell, C., Roberts, A., Goff, L., Pertea, G., Kim, D., Kelley, D.R., et al. (2012). Differential gene and transcript expression analysis of RNA-seq experiments with TopHat and Cufflinks. Nat Protoc 7(3), 562–578. doi:10.1038/nprot.2012.016.

Wang, Z., Gerstein, M., and Snyder, M. (2009). RNA-Seq: a revolutionary tool for transcriptomics. Nat Rev Genet 10(1), 57–63. doi:10.1038/nrg2484.

Yoneda, M., Guillaume, V., Sato, H., Fujita, K., Georges-Courbot, M.C., Ikeda, F., et al. (2010). The nonstructural proteins of Nipah virus play a key role in pathogenicity in experimentally infected animals. PLoS One 5(9), e12709. doi:10.1371/journal.pone.0012709.

