## Supplementary Table for "Comparison of expression profiling through microarray and RNA-seq analysis for Nipah virus"

**Supplementary Figures**

**Supplementary Figure S1.** Pie chart depicting differentially expressed downregulated genes having pathway mapped **A.** 2012 study **B.** 2017 study

**Supplementary Figure S2**. Venn diagram depicting the different and common genes among two studies

**Supplementary Figure S3.** Pie chart depicting differentially expressed upregulated genes having gene ontology based molecular function from **A.** 2012 study **B.** 2017 study

**Supplementary Figure S4.** Pie chart depicting differentially expressed downregulated genes having gene ontology based molecular function from **A.** 2012 study **B.** 2017 study

**Supplementary Figure S5.** Pie chart depicting differentially expressed downregulated genes having gene ontology based biological process from **A.** 2012 study **B.** 2017 study

**Supplementary Figure S6.** Pie chart depicting differentially expressed upregulated genes having gene ontology based cellular component from **A.** 2012 study **B.** 2017 study

**Supplementary Figure S7.** Pie chart depicting differentially expressed downregulated genes having gene ontology based cellular component from **A.** 2012 study **B.** 2017 study

**Supplementary Figure S8.** Venn diagram depicting the status of up and downregulated genes among mapped with gene ontology based molecular function in 2012 and 2017 Nipah virus studies

**Supplementary Figure S9.** Venn diagram depicting the status of up and downregulated genes among mapped with gene ontology based biological process in 2012 and 2017 Nipah virus studies

**Supplementary Figure S10.** Venn diagram depicting the status of up and downregulated genes among mapped with gene ontology based cellular component in 2012 and 2017 Nipah virus studies

**Supplementary Figure S11.** Bar graph showing the differentially expressed downregulated genes having GAD disease category from **A.** 2012 study **B.** 2017 study

**Supplementary Figure S12.** The signal transduction pathway mapped of the upregulated genes from 2012 study using reactome software

**Supplementary Figure S13.** The metabolism pathway mapped of the upregulated genes from 2012 study using reactome software

**Supplementary Figure S14.** Pathway mapped of the down regulated genes from 2012 study using reactome software

**Supplementary Figure S15.** The immune pathway mapped of the downregulated genes from 2012 study using reactome software

**Supplementary Figure S16.** The signal transduction pathway mapped of the downregulated genes from 2012 study using reactome software

**Supplementary Figure S17.** The metabolism pathway mapped of the upregulated genes from 2017 study using reactome software

**Supplementary Figure S18.** The signal transduction pathway mapped of the upregulated genes from 2017 study using reactome software

**Supplementary Figure S19.** Pathway mapped of the downregulated genes from 2017 study using reactome software

**Supplementary Figure S20.** The signal transduction pathway mapped of the downregulated genes from 2017 study using reactome software

**Supplementary Figure S21.** The metabolism pathway mapped of the downregulated genes from 2017 study using reactome software

**Supplementary Figure S22.** Rank-Rank scatter plot between 2012 and 2017 studies

**Supplementary Figure S23.** Venn diagram depicting the up and down regulate genes (differentially expressed genes) among 2012 and 2017 studies

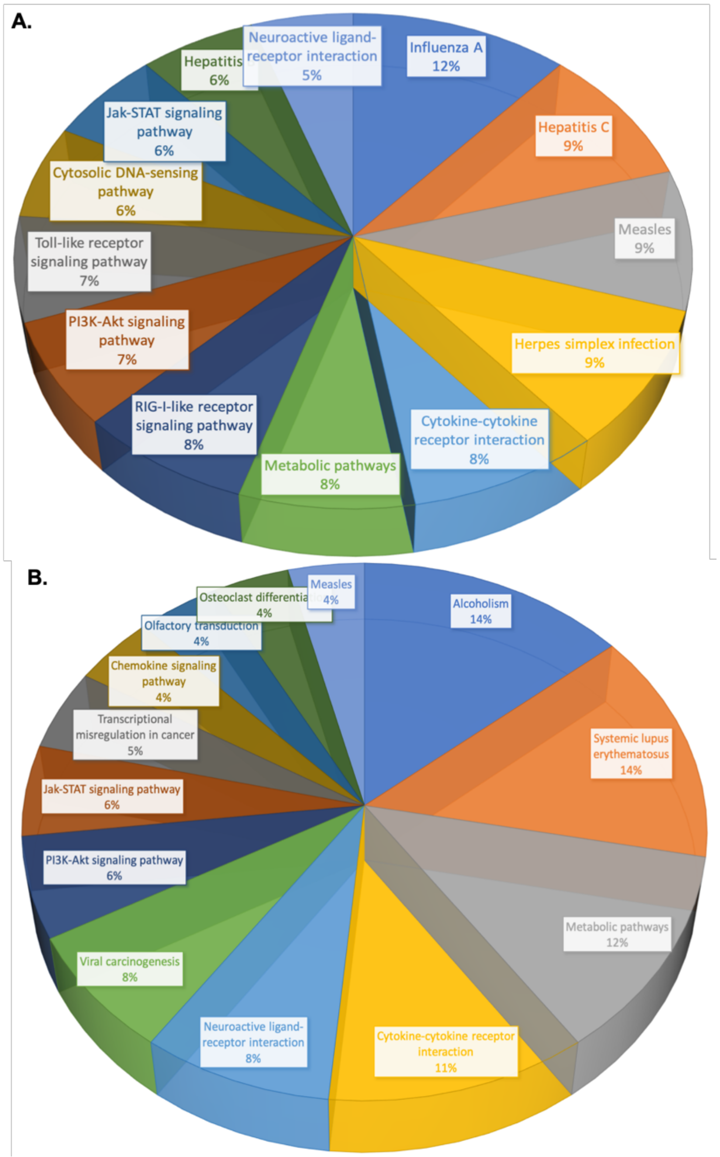


**Supplementary Figure S1.** Pie chart depicting differentially expressed downregulated genes having pathway mapped **A.** 2012 study **B.** 2017 study

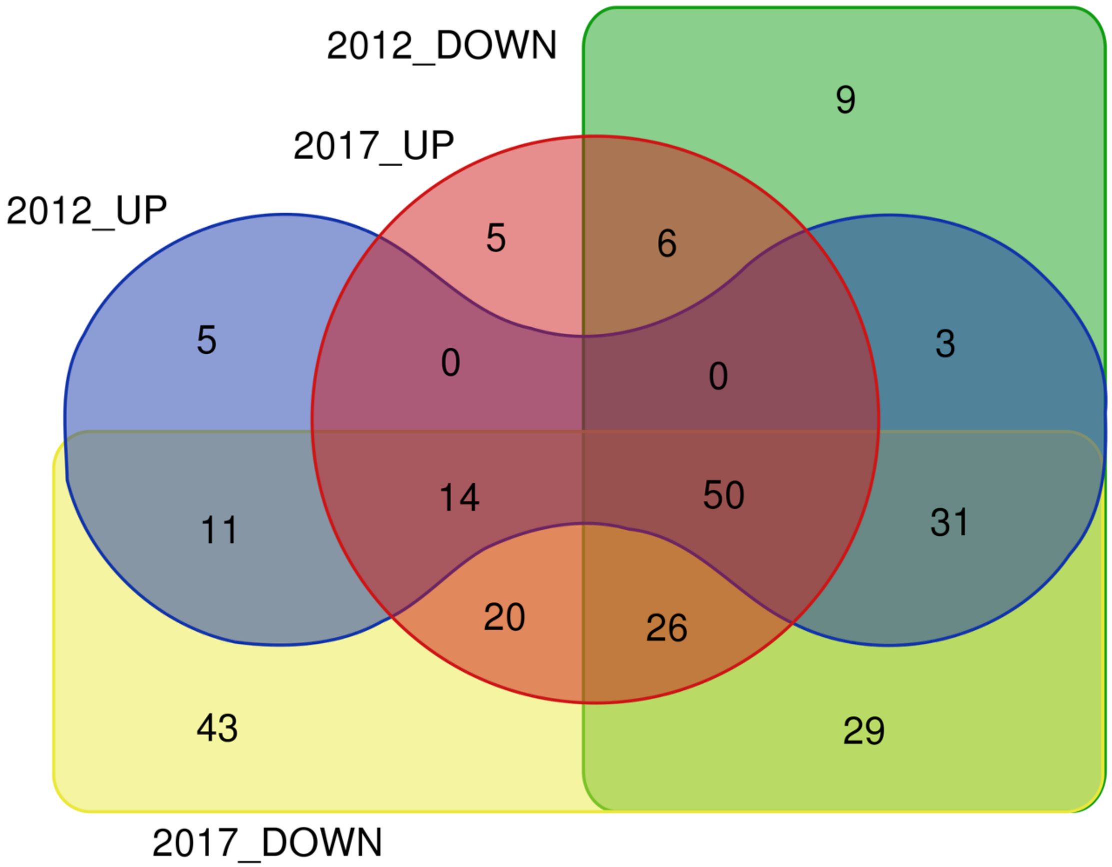


**Supplementary Figure S2**. Venn diagram depicting the different and common genes among two studies

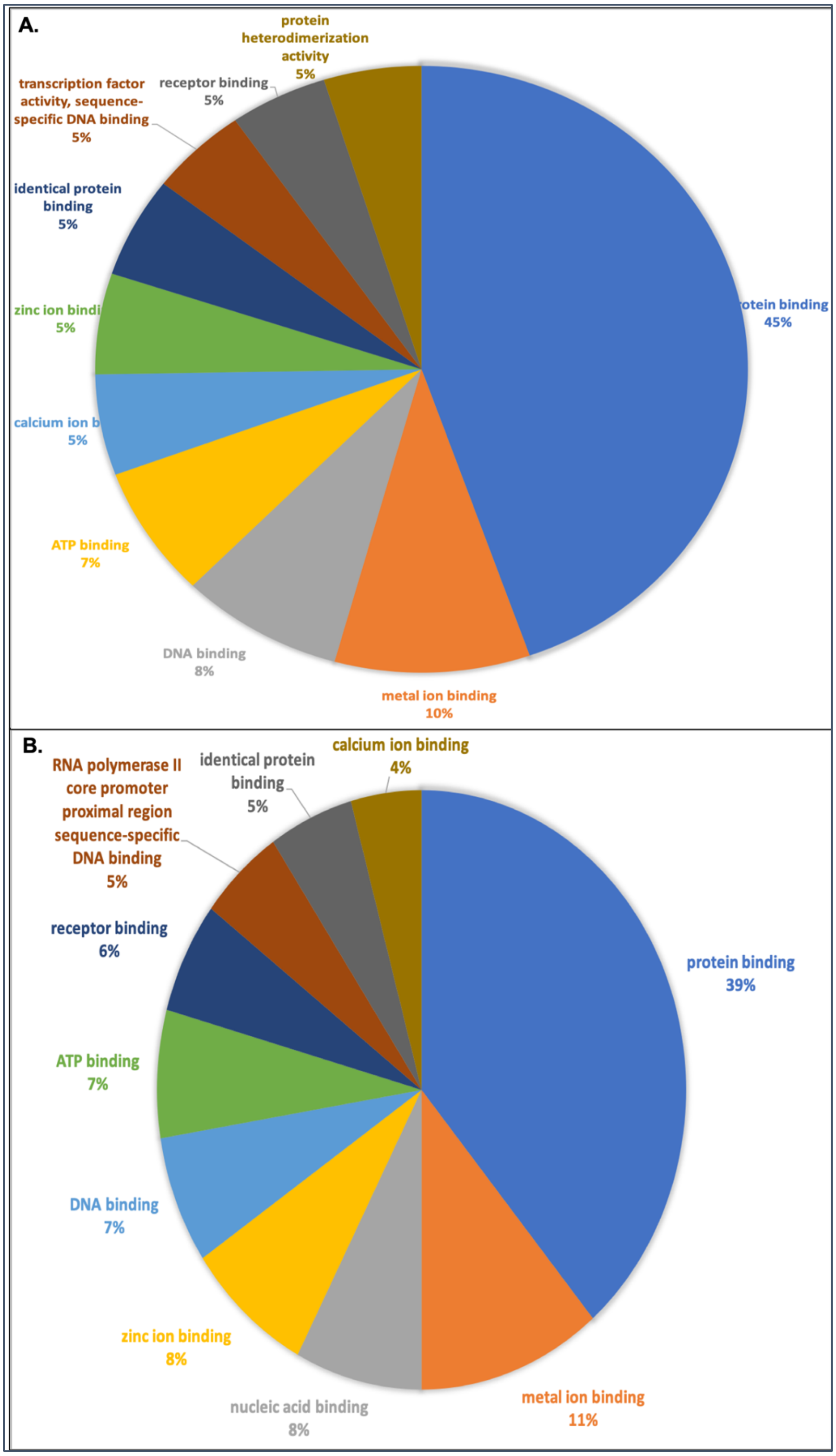


**Supplementary Figure S3.** Pie chart depicting differentially expressed upregulated genes having gene ontology based molecular function from **A.** 2012 study **B.** 2017 study

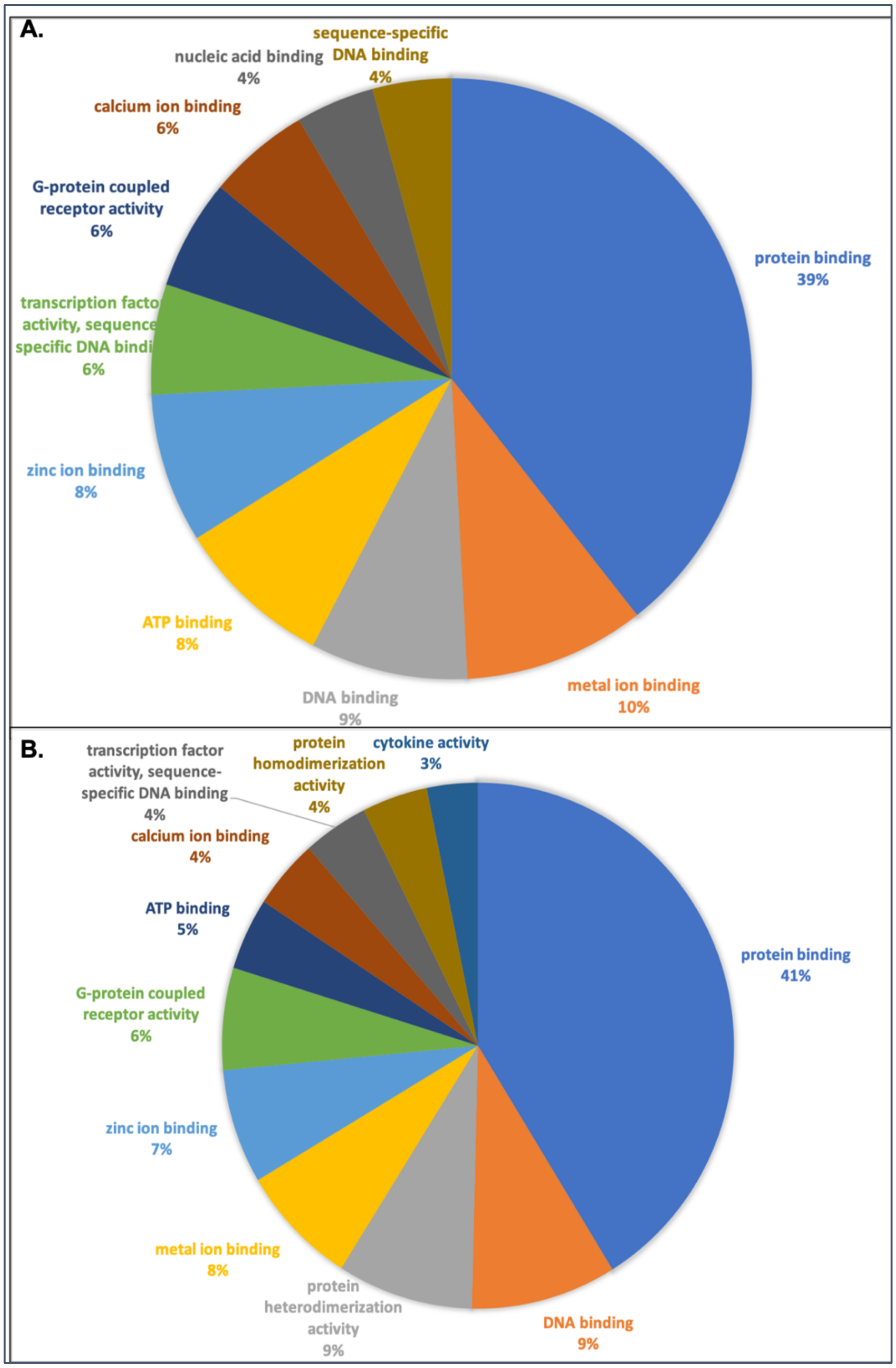


**Supplementary Figure S4.** Pie chart depicting differentially expressed downregulated genes having gene ontology based molecular function from **A.** 2012 study **B.** 2017 study

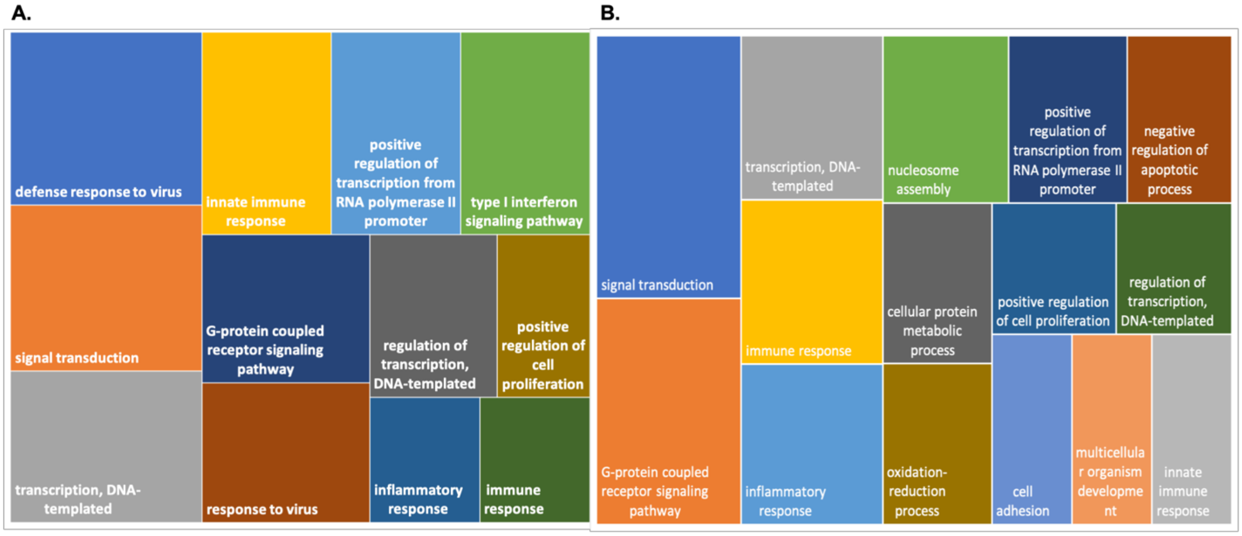


**Supplementary Figure S5.** Pie chart depicting differentially expressed downregulated genes having gene ontology based biological process from **A.** 2012 study **B.** 2017 study

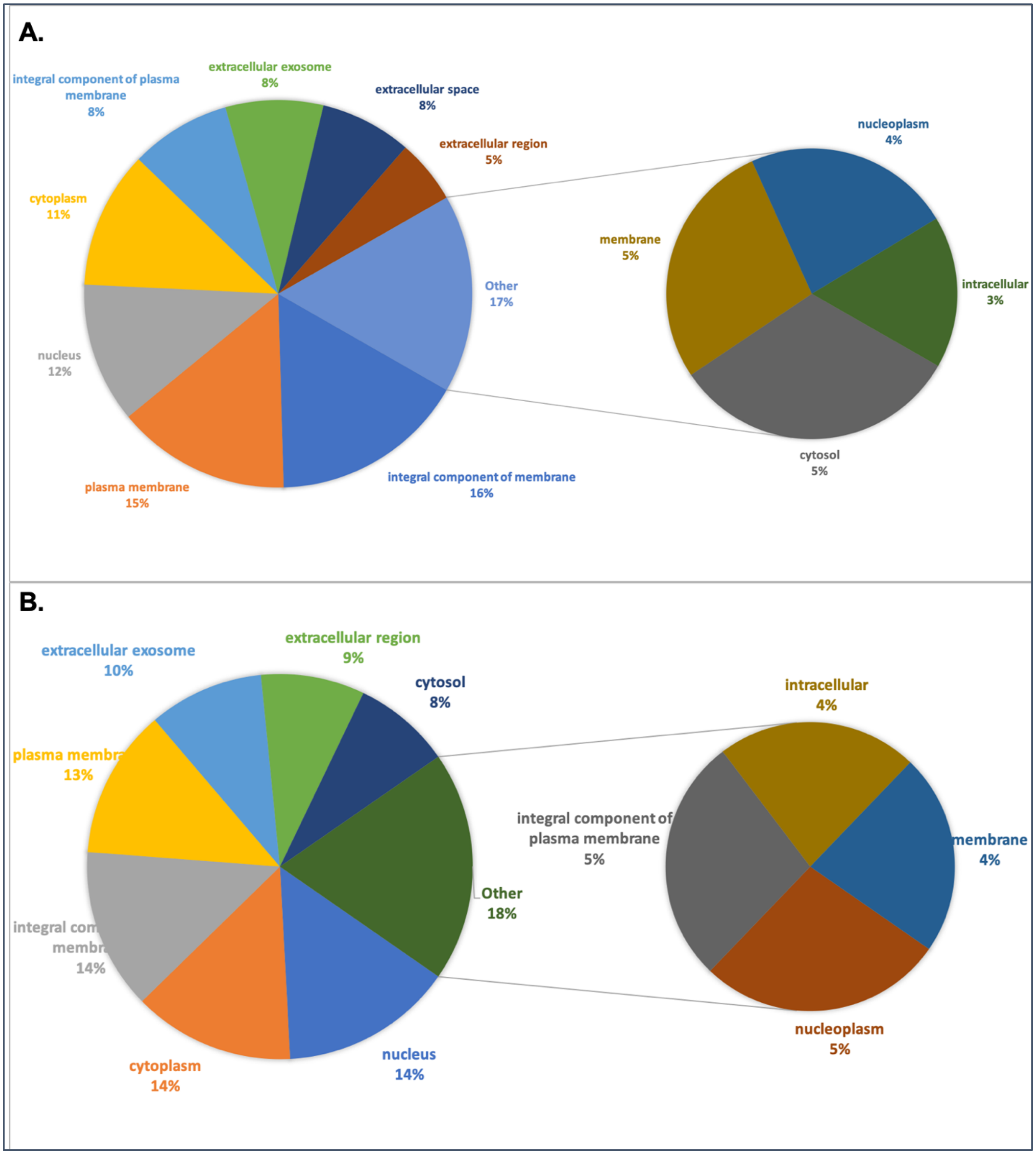


**Supplementary Figure S6.** Pie chart depicting differentially expressed upregulated genes having gene ontology based cellular component from **A.** 2012 study **B.** 2017 study

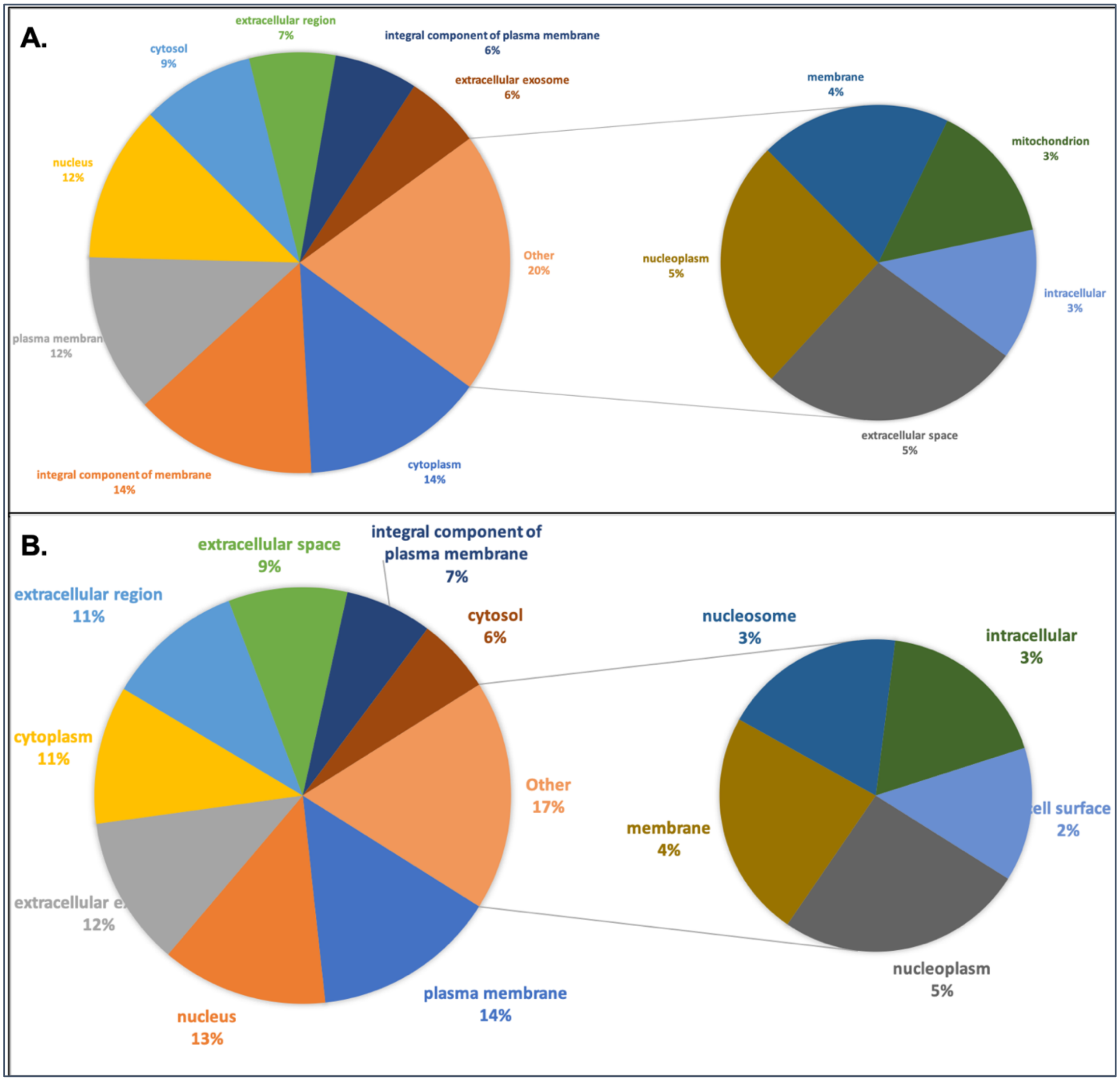


**Supplementary Figure S7.** Pie chart depicting differentially expressed downregulated genes having gene ontology based cellular component from **A.** 2012 study **B.** 2017 study

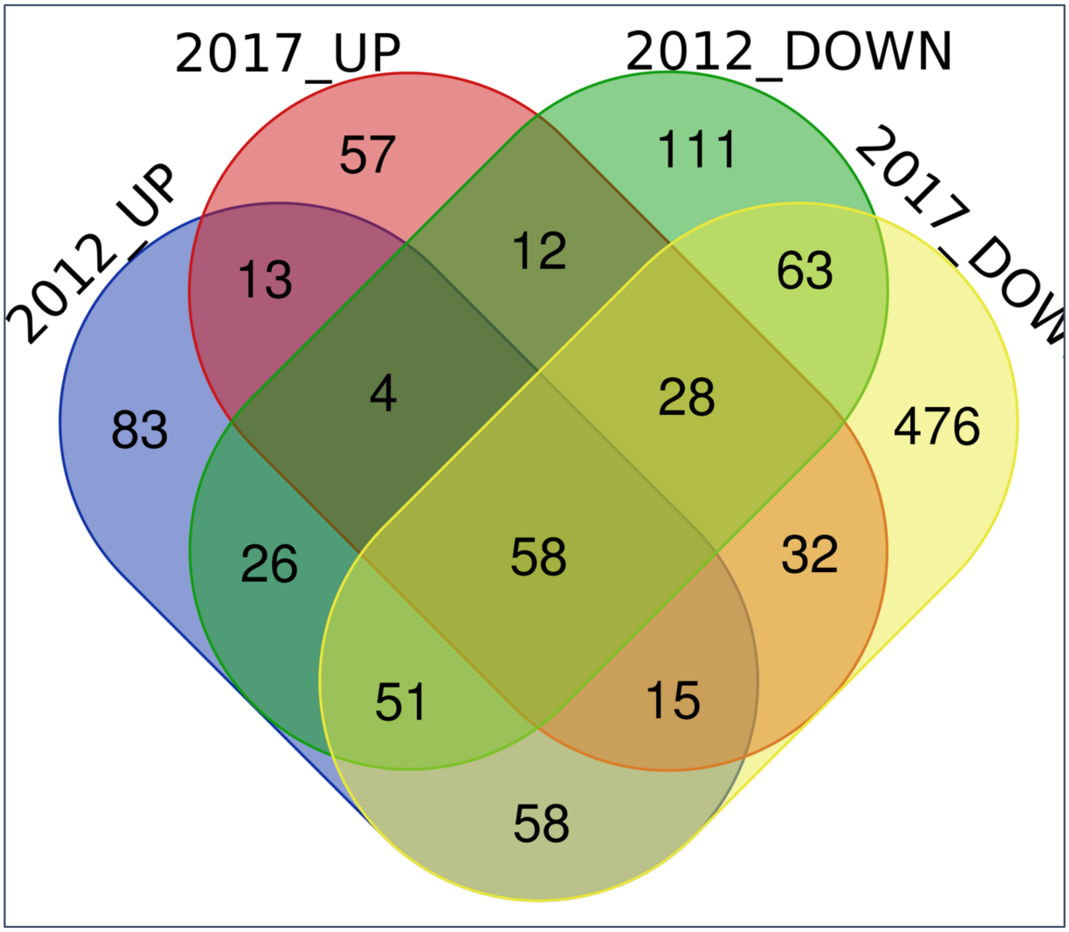


**Supplementary Figure S8.** Venn diagram depicting the status of up and downregulated genes among mapped with gene ontology based molecular function in 2012 and 2017 Nipah virus studies

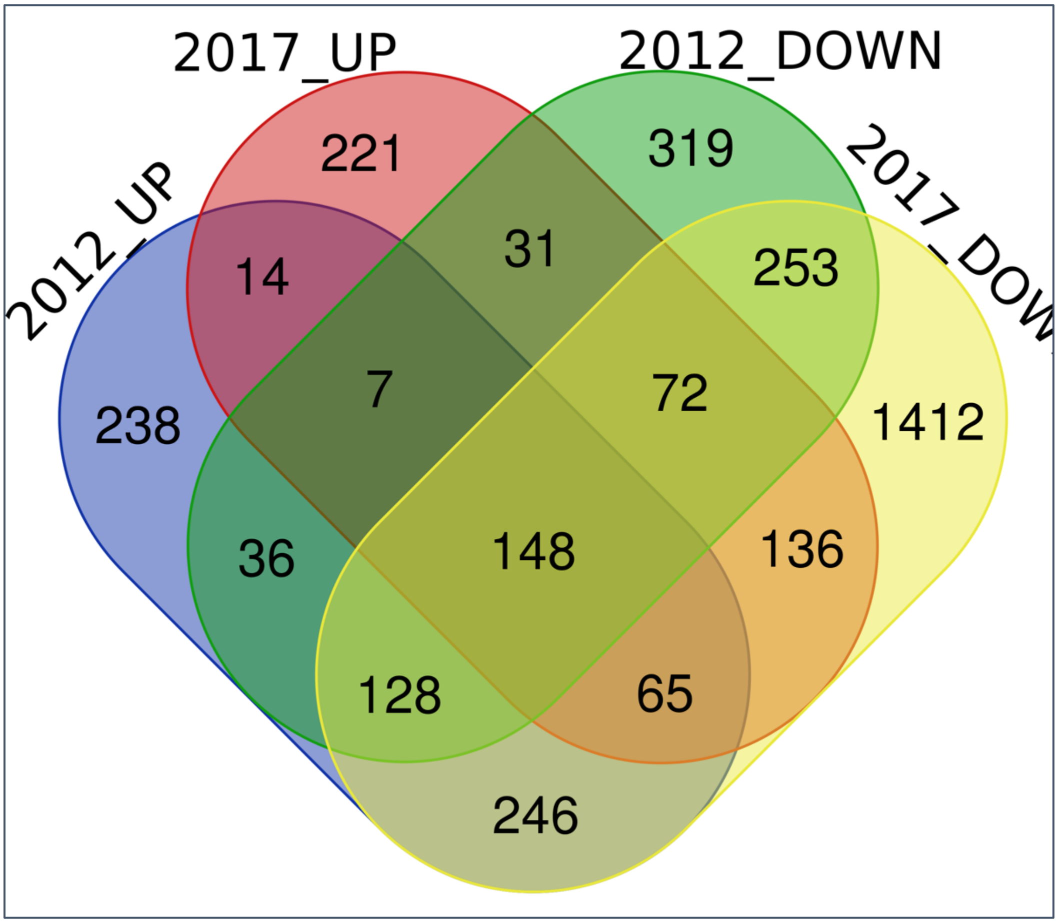


**Supplementary Figure S9.** Venn diagram depicting the status of up and downregulated genes among mapped with gene ontology based biological process in 2012 and 2017 Nipah virus studies

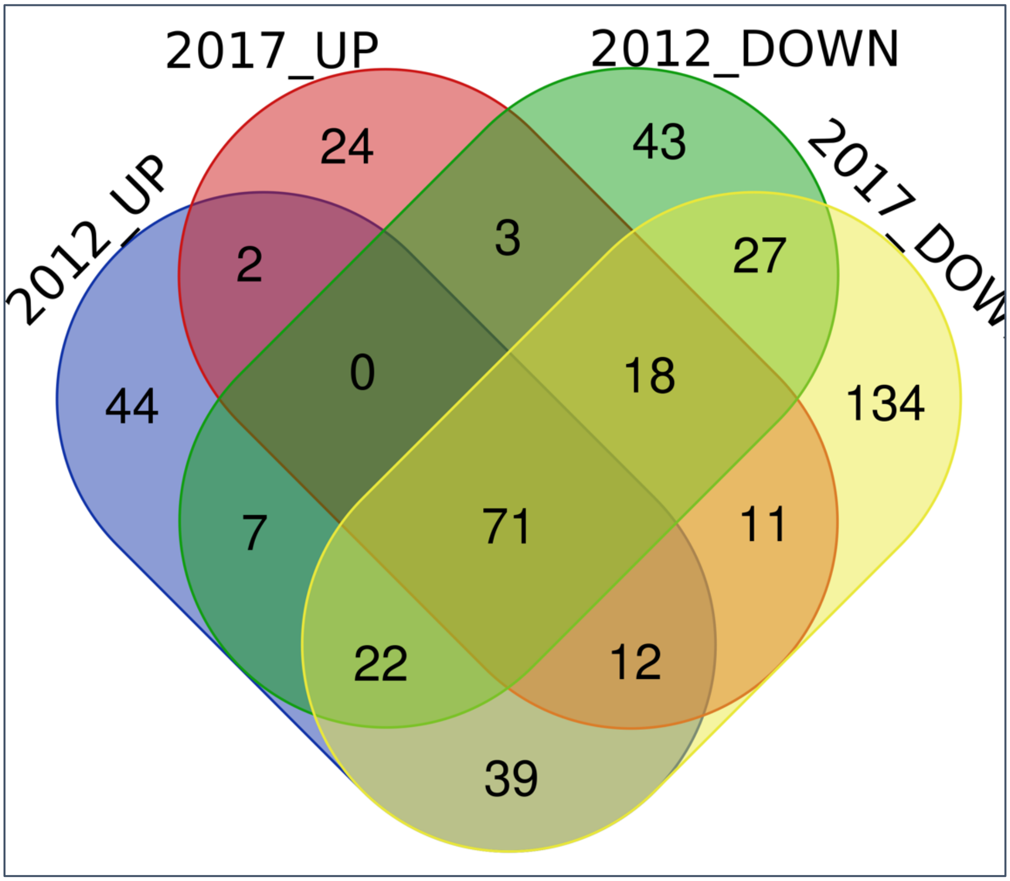


**Supplementary Figure S10.** Venn diagram depicting the status of up and downregulated genes among mapped with gene ontology based cellular component in 2012 and 2017 Nipah virus studies

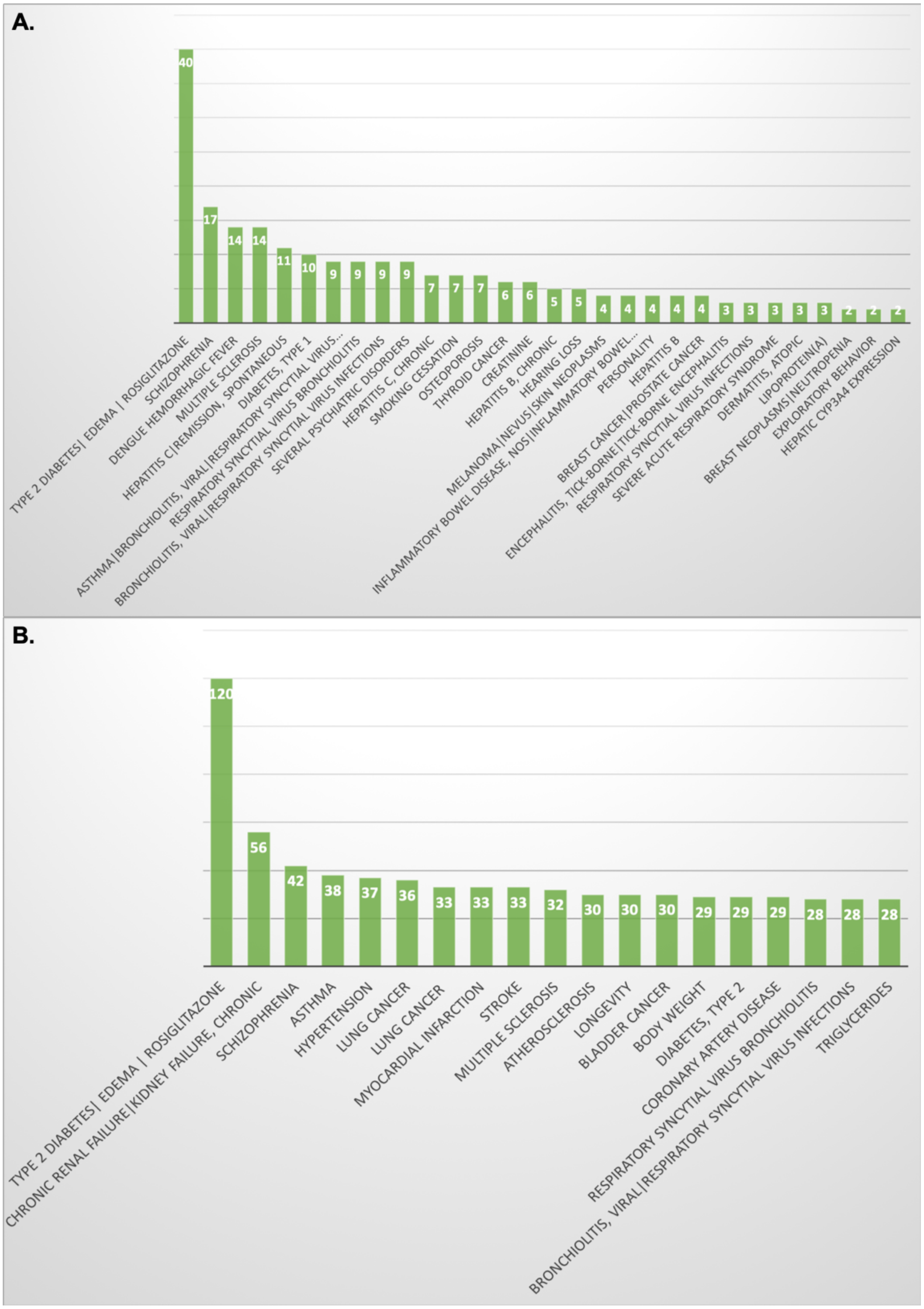


**Supplementary Figure S11.** Bar graph showing the differentially expressed downregulated genes having GAD disease category from **A.** 2012 study **B.** 2017 study

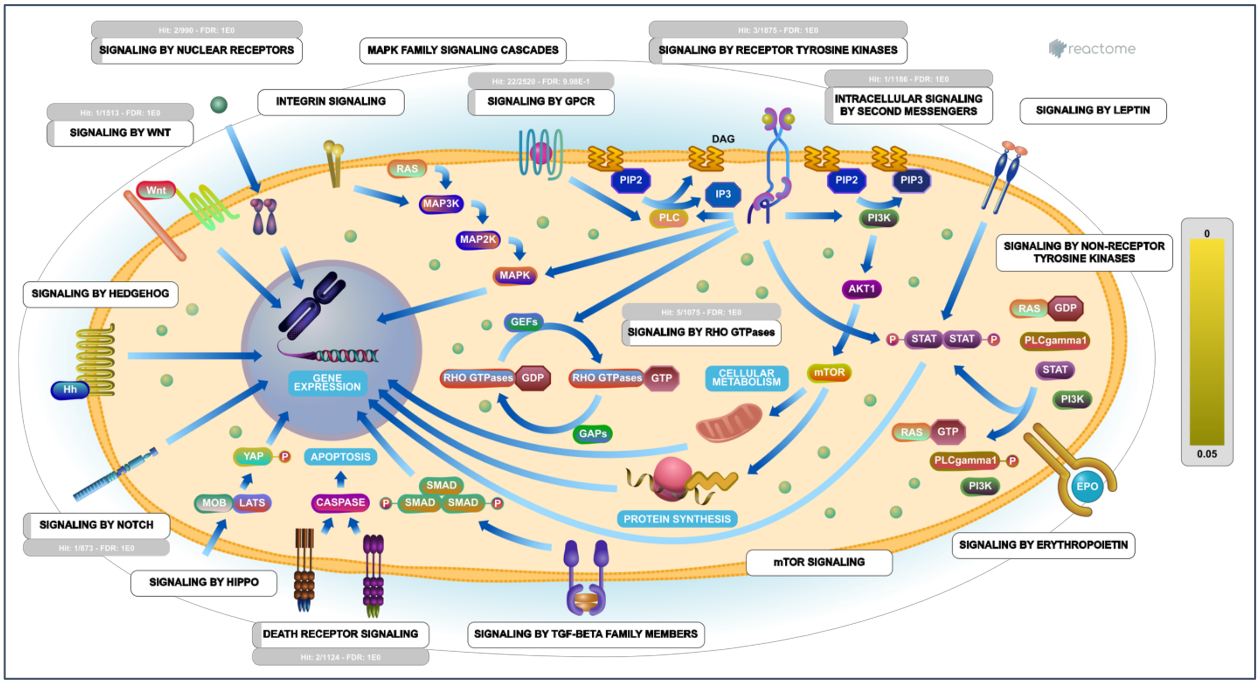


**Supplementary Figure S12.** The signal transduction pathway mapped of the upregulated genes from 2012 study using reactome software

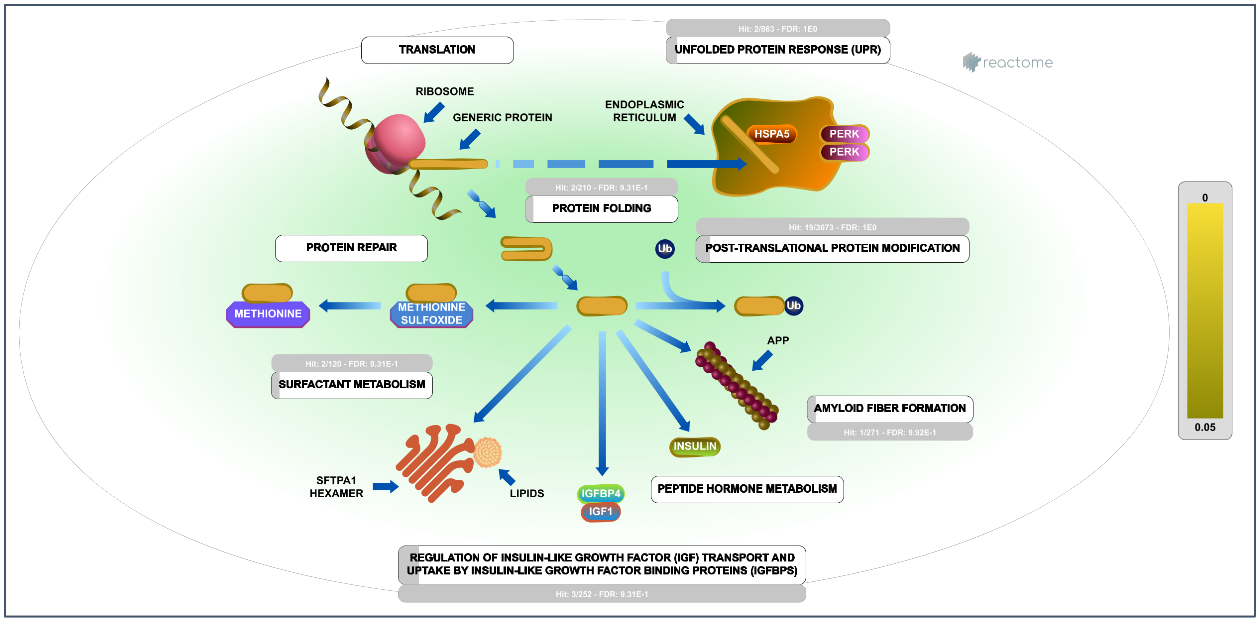


**Supplementary Figure S13.** The metabolism pathway mapped of the upregulated genes from 2012 study using reactome software

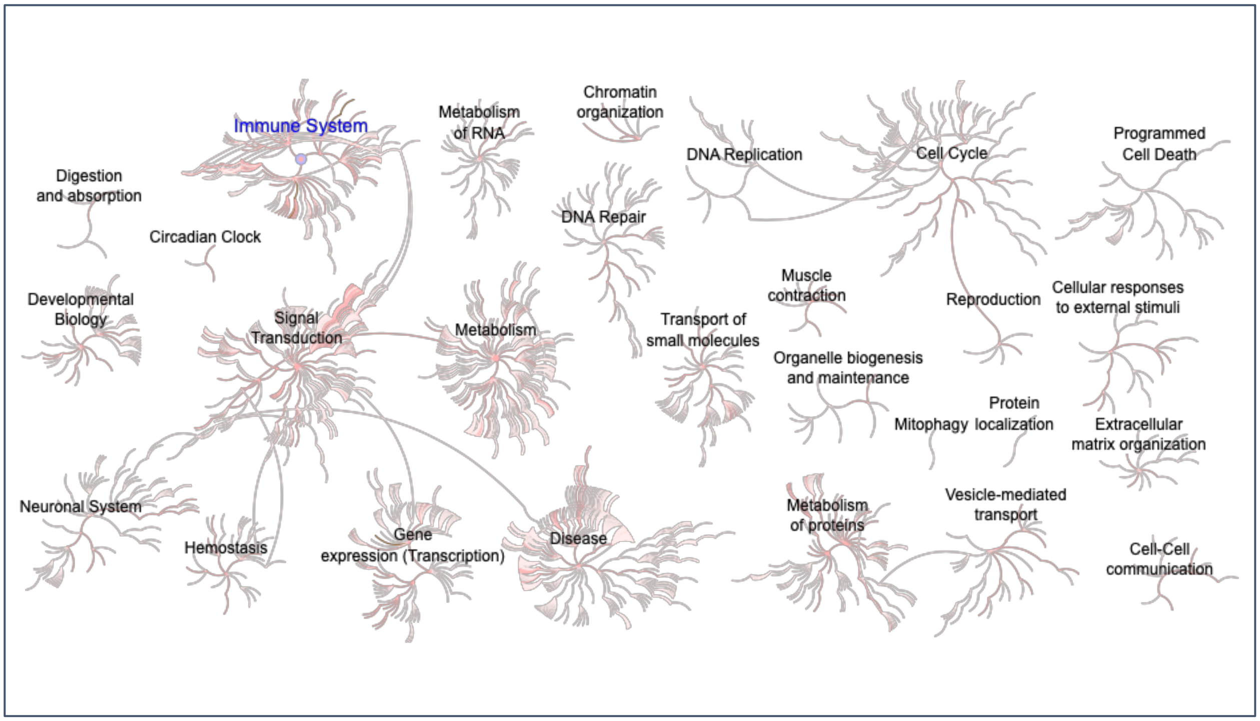


**Supplementary Figure S14.** Pathway mapped of the down regulated genes from 2012 study using reactome software

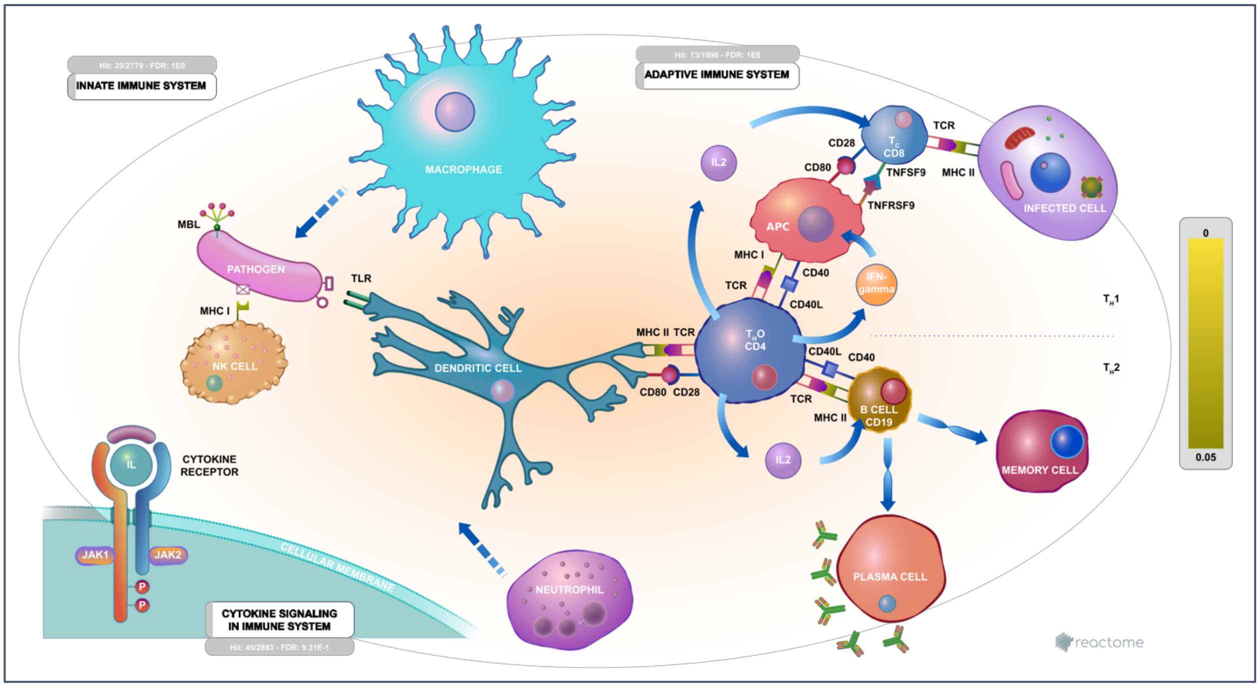


**Supplementary Figure S15.** The immune pathway mapped of the downregulated genes from 2012 study using reactome software

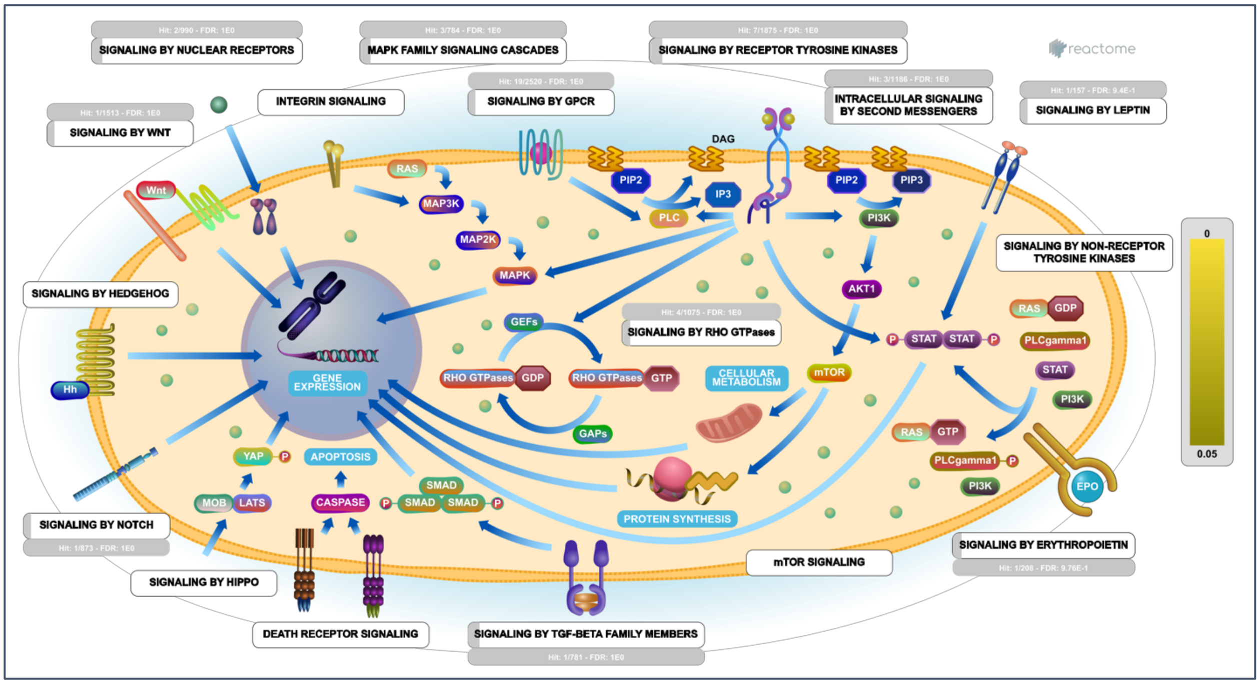


**Supplementary Figure S16.** The signal transduction pathway mapped of the downregulated genes from 2012 study using reactome software

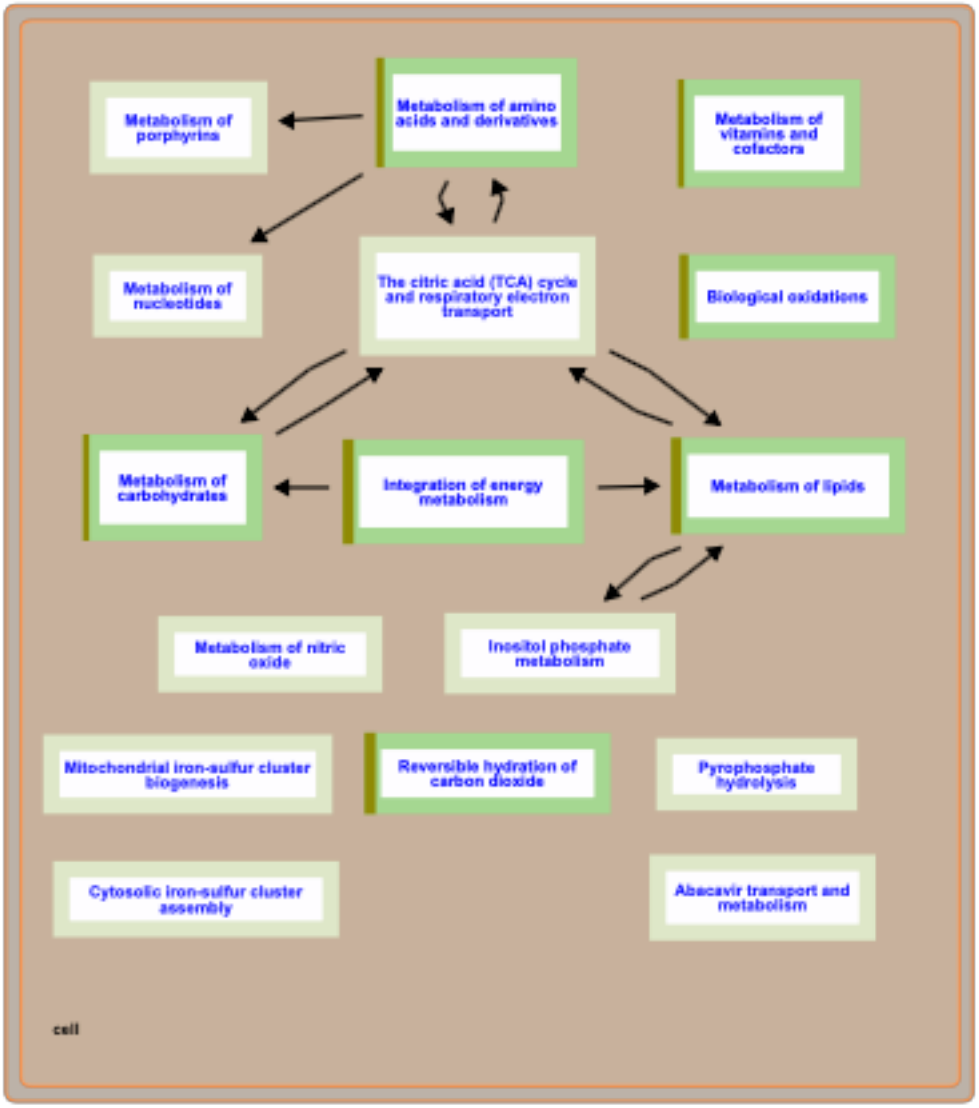


**Supplementary Figure S17.** The metabolism pathway mapped of the upregulated genes from 2017 study using reactome software

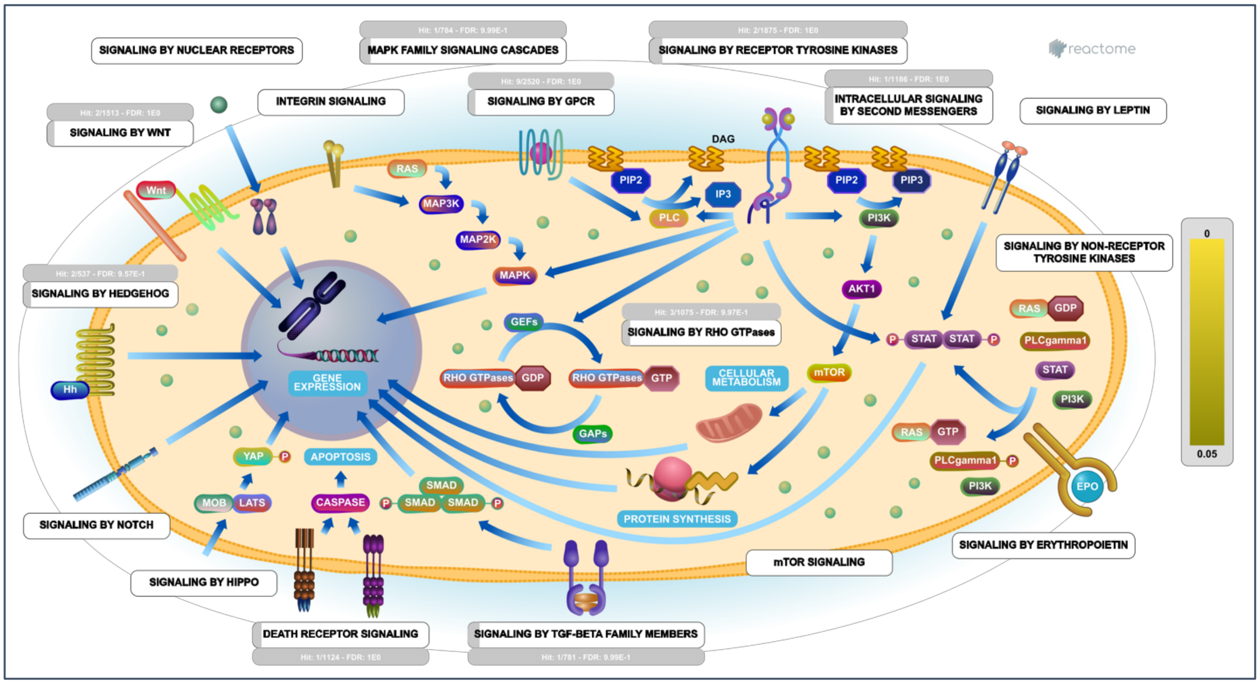


**Supplementary Figure S18.** The signal transduction pathway mapped of the upregulated genes from 2017 study using reactome software

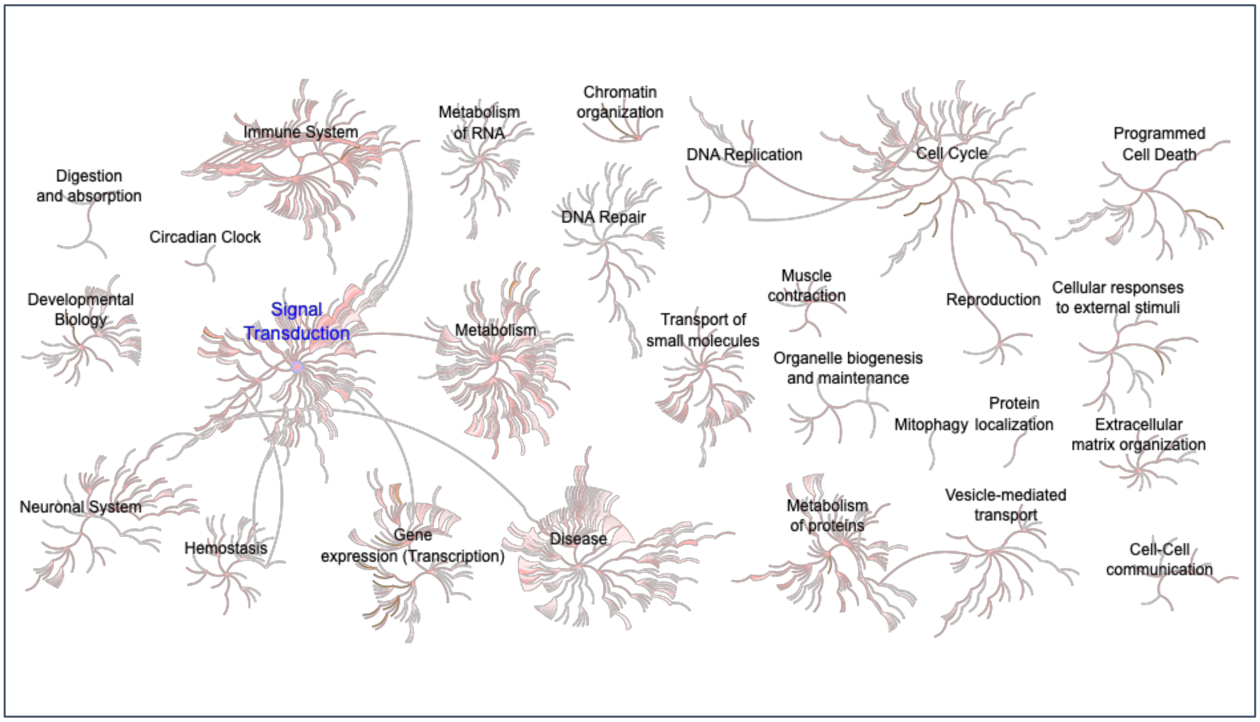


**Supplementary Figure S19.** Pathway mapped of the downregulated genes from 2017 study using reactome software

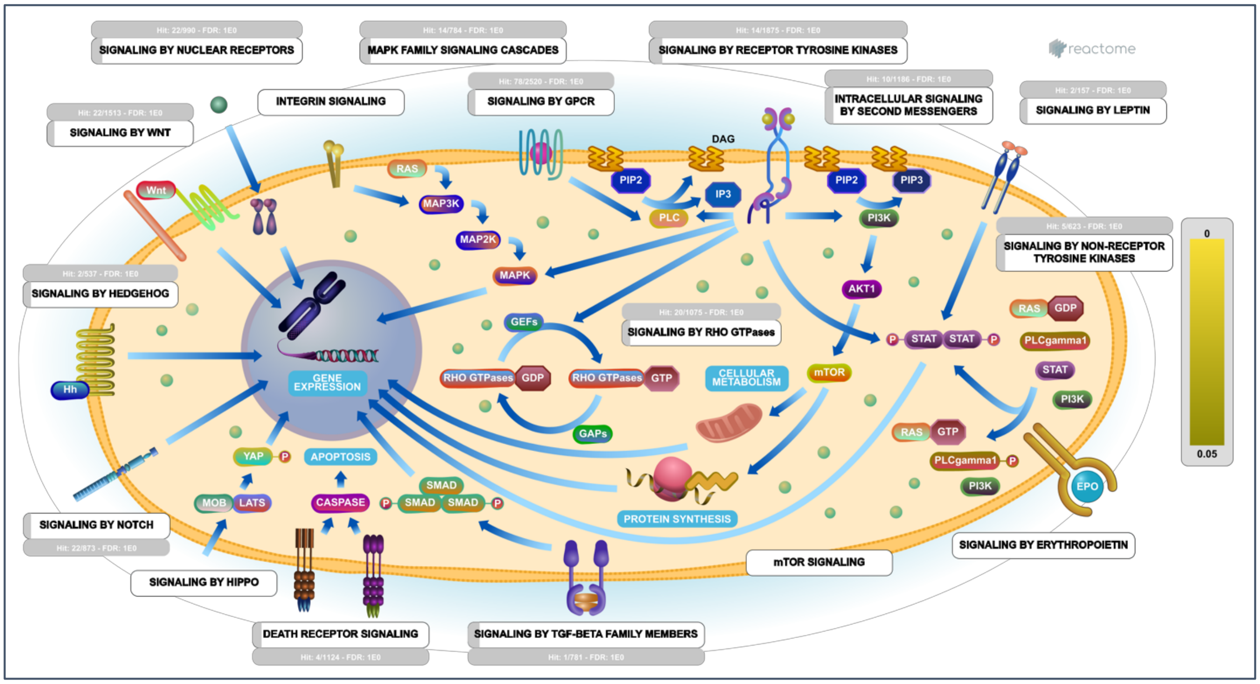


**Supplementary Figure S20.** The signal transduction pathway mapped of the downregulated genes from 2017 study using reactome software

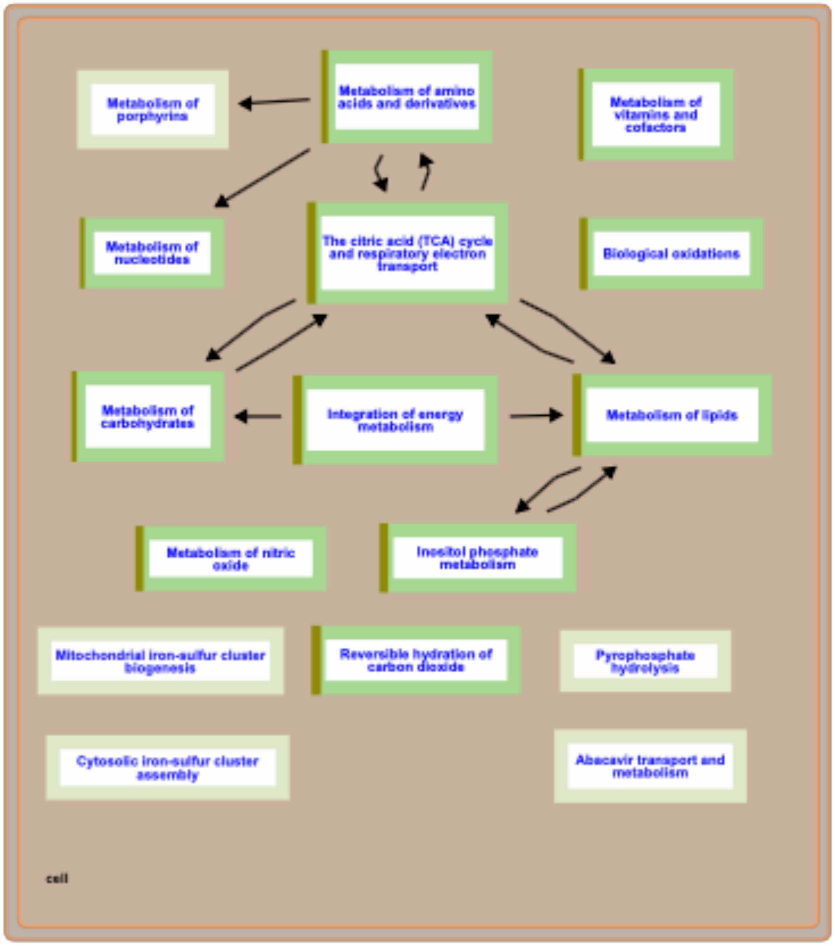


**Supplementary Figure S21.** The metabolism pathway mapped of the downregulated genes from 2017 study using reactome software

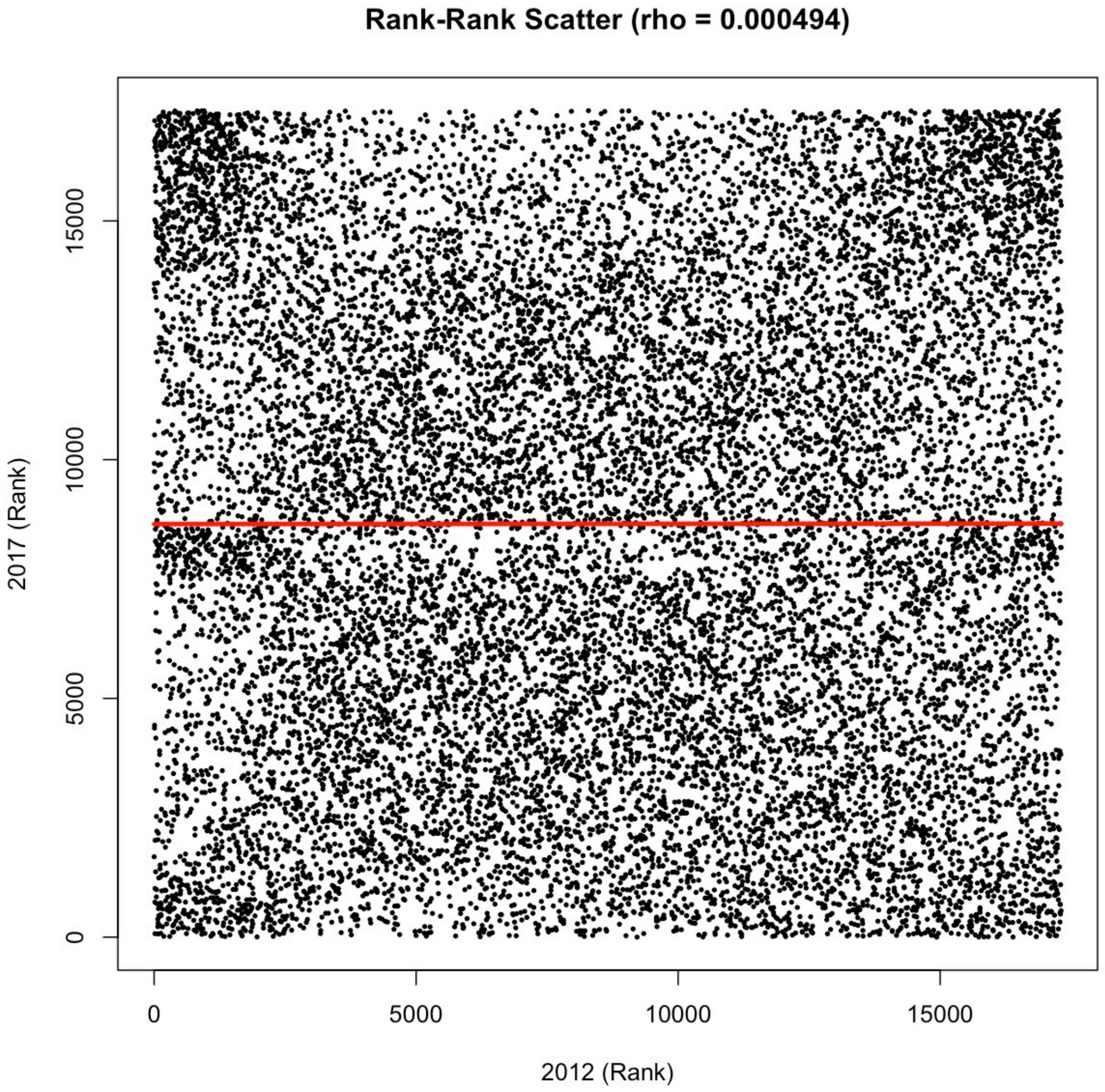


**Supplementary Figure S22.** Rank-Rank scatter plot between 2012 and 2017 studies

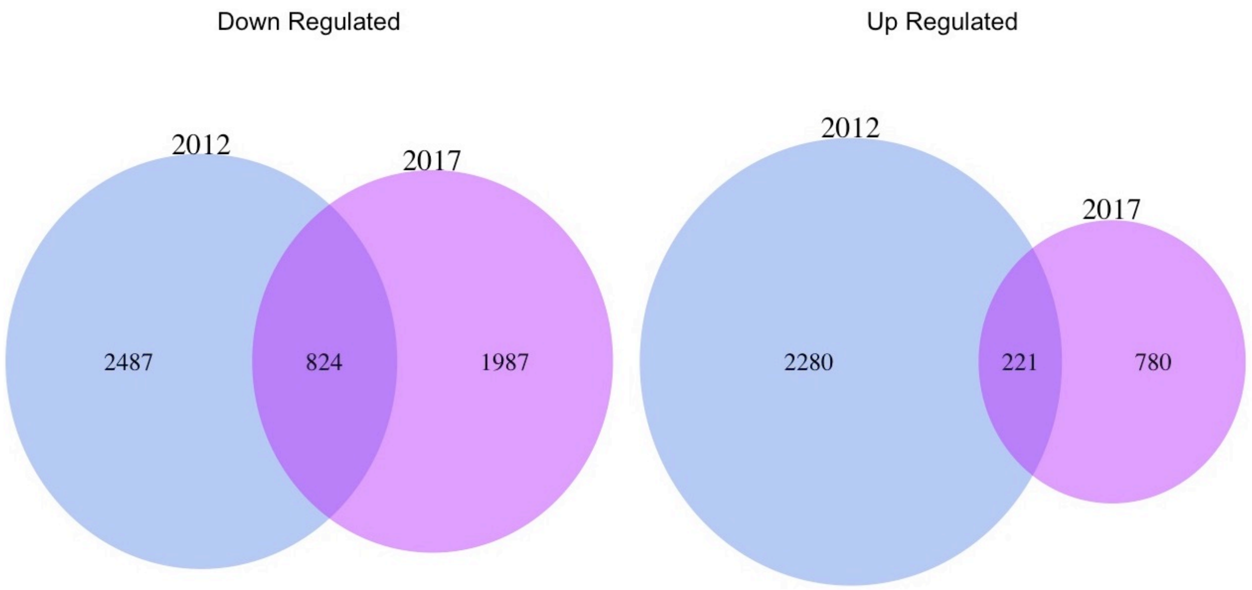


**Supplementary Figure S23.** Venn diagram depicting the up and down regulate genes (differentially expressed genes) among 2012 and 2017 studies
